# Dynamism in paintings and photographs: how the power of diagonals in art overcomes the oblique effect in visual perception

**DOI:** 10.64898/2026.07.22.739875

**Authors:** Serena Castellotti, Elvio Blini, Maria Del Viva, Qasim Zaidi

**Author notes:** Corresponding author: Qasim Zaidi; Complete address: Graduate Center for Vision Research, State University of New York College of Optometry, 33 West 42nd Street, New York, NY 10036, USA. **Author Contributions:** QZ conceived the research question. SC, MDV, and QZ designed Experiments 1 and 2. SC and EB programmed and conducted the experiments, and SC and QZ analyzed the data. SC and MDV designed Experiment 3, while SC and EB conducted the experiment and performed the analyses. SC and QZ wrote the manuscript, with substantial input from EB and MDV.

## Abstract

Artists and photographers use diagonals to convey dynamism, tension and instability in naturalistic and abstract images. However, the oblique effect in visual perception refers to a reduced sensitivity to oblique compared to cardinal (vertical/horizontal) orientations, linked to anisotropy in the distribution of preferred orientations of primary visual cortex neurons, which reflect the statistical distribution of edge orientations in natural images. Distal diagonals generically project onto oblique orientations on the retina, creating a conundrum between their perceptual salience in art and their reduced sensitivity in vision.

We conjectured that global spatial configurations could shape perceived dynamism overriding local orientations. Inspired by van Doesburg’s 1929 Arithmetical Composition, we created 60 distinct linear configurations of four identically-shaped rhombi each, in which global and local-edge orientations were independently manipulated to be cardinal or oblique, and measured their perceived dynamism with behavioral, psychophysical, and psychophysiological methods.

First, 250 observers rated oblique configurations as more dynamic, tridimensional, and unstable than cardinal configurations, with wedge shapes and element spacing enhancing dynamism. Second, pairwise comparisons (50 observers) yielded a perceptual dynamism scale that confirmed the dominant role of global orientations. Third, pupillometry (35 observers) showed that pupil dilation correlates systematically with subjective dynamism ratings.

Perceived dynamism in static images is thus driven primarily by the orientation and shape of spatial configurations, indicating the importance of estimating orientations and shapes of multi-element objects beyond decoding local orientations. By linking visual structure to emotional perceptual experience, this investigation provides empirically grounded principles for creating dynamic percepts in visual art.

**SIGNIFICANCE STATEMENT:** Diagonals have been used since Michelangelo to create dynamic compositions in representative and abstract art and photography, but paradoxically humans have been shown to be less sensitive to oblique orientations like those projected on the retina by diagonal edges.

We resolve this conflict by distinguishing between global configurations versus local edges and showing through behavioral, psychophysical, and psychophysiological experiments that the global orientation and shape of a multi-element configuration has a much larger effect on perceived dynamism than do the edge orientations of its elements.

Neural estimation of orientations and shapes of multi-element configurations is thus essential for image understanding. Our results provide perceptually grounded bases for using diagonal configurations with wedge shapes to create dynamic percepts in visual art.

## INTRODUCTION

Art and photography classes teach about the power of diagonals. On the other hand, the oblique effect in visual perception refers to a reduced sensitivity to oblique orientations compared to the cardinal horizontal and vertical orientations (1), which is probably due to the greater number and orientation selectivity of neurons tuned to cardinal orientations in the primary visual cortex (2, 3). Since diagonals in images project to oblique orientations on the retinae, this creates a conundrum between the saliency of diagonals in art against the weakness of oblique orientations in perception. We describe the conundrum in detail and then resolve it with empirical experiments.

**Fig. 1A** shows Michelangelo’s Crucifixion of St Peter fresco (c. 1546–1550) and **Fig. 1B** the preparatory drawing. Scully (4) claimed that it was the first diagonal composition in painting and described it dramatically: “Space moves diagonally in one mighty reflex, extending itself psychically far beyond the confines of the wall”. The dynamism of the whole scene is enhanced by the diagonal cross, but even the isolated drawing of the diagonal figure is dynamic, seemingly falling in space. The dynamism of diagonals versus verticals is further illustrated by comparing Caravaggio’s diagonal Crucifixion of Saint Peter (1601; **Fig. 1C**) to Guido Reni’s vertical version (1604; **Fig. 1D**) commissioned for S. Paolo alle Tre Fontane, clearly showing Caravaggio’s influence but with lessened dramatic tension.

**Fig. 1.**
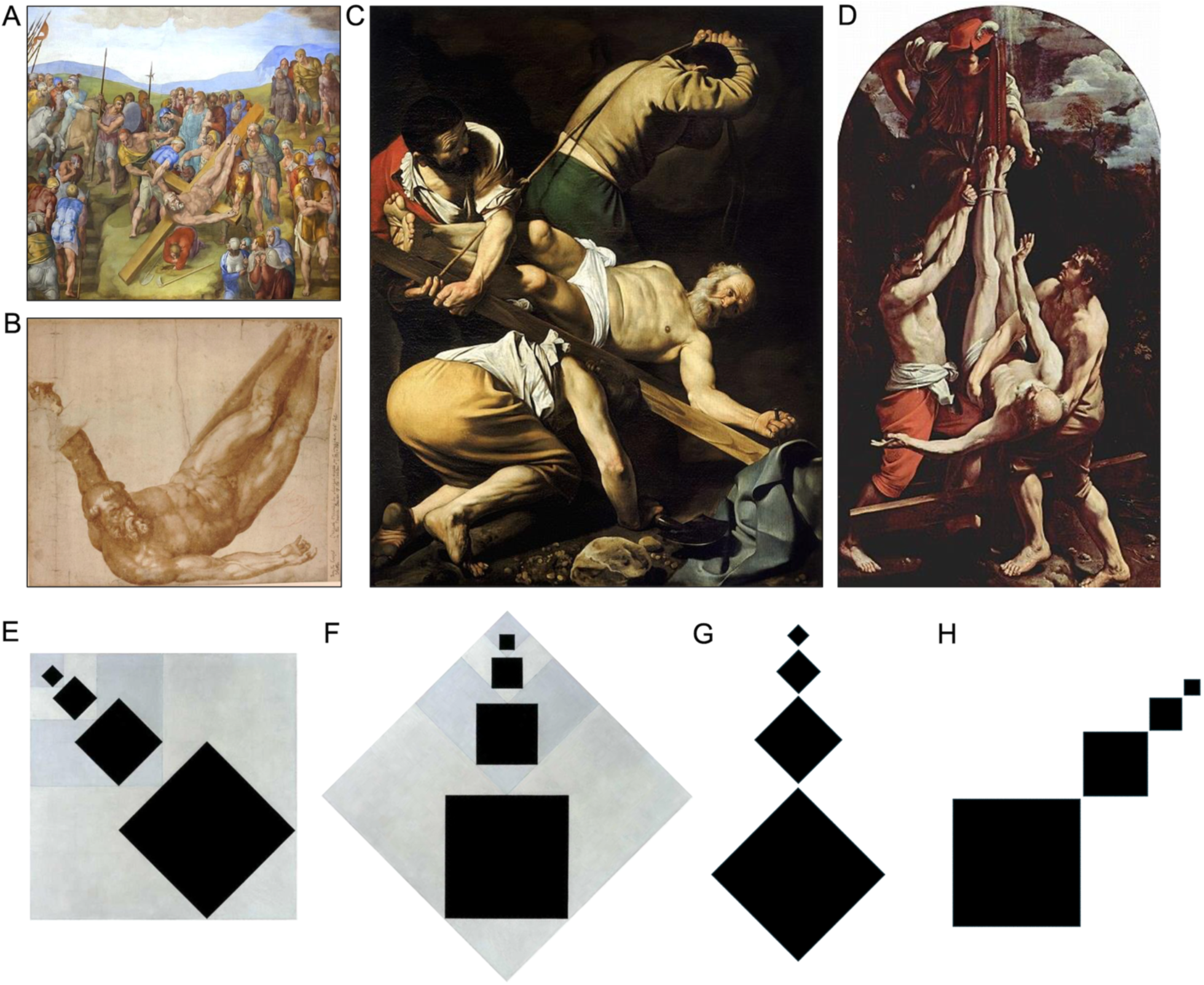
From diagonality in pictorial compositions to experimental stimulus design. **(A)** Michelangelo’s Crucifixion of St Peter (c. 1546–1550) in the Vatican Cappella Paolina, Rome **(B)** Preparatory drawing for the fresco**. (C)** Caravaggio’s Crucifixion of Saint Peter (1601) in Santa Maria del Popolo, Rome**. (D)** Guido Reni’s Crucifixion of Saint Peter (1604) in the Pinacoteca Vaticana, Rome. **(E)** Theo van Doesburg’s Arithmetical Composition (1929–1930), characterized by both oblique local edges and a diagonal global configuration. **(F)** A version of the same composition rotated by 45° counterclockwise, resulting in cardinal local edges and a cardinal global configuration. **(G)** Example experimental stimulus with a cardinal global configuration and oblique local edges. **(H)** Example experimental stimulus with a diagonal global configuration and cardinal local edges. All paintings are in the public domain.

The dynamic contributions of diagonals extend to abstract painting, and Kandinsky, who experimented with the elements of representation, said that diagonals “can be looked upon as a kind of measure of tension” (5). He reasoned that something diagonal is generally unstable or already falling. A cursory web search for diagonals in photography finds several quotes such as “Compositions with domination of horizontal and vertical lines are architectonical and static. In comparison with them compositions comprised of diagonal lines are dynamic” (6), “Diagonals are always dynamic elements within a picture. They create a strong feeling of movement because the eye wants to move along a diagonal” (7) and “The most dynamic compositions are the ones based on diagonal lines and foreshortenings” (8). These quotes are probably based on extensive experience, so worth examining empirically and conceptually.

In visual neuroscience, an analysis of orientation processing by 4,418 neurons in the cat’s primary visual cortex showed the largest number preferentially tuned to horizontal, then vertical, and then oblique, whereas orientation tuning width was narrowest for horizontally tuned cells, and narrower for vertically tuned cells than the neighboring oblique orientations (3). In addition, optical imaging analysis of 12 years of responses from the lower visual field of 79 hemispheres of 58 adult macaque monkeys to four orientations of gratings showed clear orientation anisotropies in macaque V1 and V4 (9). For V1, orientation anisotropy was both visual field location-independent (cardinal) and location-dependent (radial), while V4 showed the largest number of neurons preferentially tuned to horizontal, then vertical, and then oblique. Moreover, in humans, the fMRI BOLD response is greater to cardinal stimuli than to oblique stimuli in V1, but extrastriate visual areas (V2, V3, VP) did not show a reliable oblique effect (2). Anisotropic neuronal distributions are thought to underlie the anisotropies in psychophysical orientation estimation and discrimination, which are worse for oblique orientations (1), with V1 BOLD response amplitudes well correlated with both contrast detection and orientation discrimination (2). The anisotropy in neuronal distributions matches the local orientation distribution measured in photographs of natural scenes, suggesting that anisotropic neural population responses are the substrate for observers’ internal models for orientation (10), which is also supported by models for the failures of 2D and 3D shape constancy under rotation in the image plane (11, 12).

To explain why diagonals are dynamic despite the possible deficiency in neuronal processing of oblique orientations, we examined the claim of dynamism empirically while testing for static cues that promote percepts of dynamism in still images. We considered several ideas. First, novelty detection may be enhanced by adaptation to stimulus statistics, making the more frequent cardinal (horizontal and vertical) orientations less salient and potentially causing oblique edges or contours to pop out. Second, when most local orientations in an image are cardinal, the implicit frame they form can further highlight oblique features. Third, extended diagonal forms can generate percepts of depth or tridimensionality, allowing figures to appear as if they are emerging out of the picture plane. Fourth, real-world objects that are neither vertical nor horizontal are often in motion – falling or moving intentionally – so diagonal configurations may carry an intrinsic dynamic meaning. The first two ideas concern orientations of local edges/contours, while the latter two concern the global orientation of extended configurations, raising the critical question of whether diagonal configurations create greater dynamic effects than diagonal edges/contours.

The stimuli we constructed were inspired by Theo van Doesburg’s painting Arithmetical Composition (1929–1930; **Fig. 1E**). Doesburg and Mondrian founded the De Stijl movement, which pursued pure abstraction through compositions built only from horizontal and vertical grids, but when Doesburg began introducing diagonals into his work, this departure from cardinality was powerful enough to cause an irreparable rift. The original painting (**Fig. 1E**) illustrates the dynamism of diagonals, especially when compared with the rotated version in **Fig. 1F**. However, local edges and global configurations are both oblique in **Fig. 1E** and cardinal in **Fig. 1F**, so their relative contributions cannot be inferred from this comparison. For that, we created the variations in **Fig. 1G** with oblique local edges but a vertical global configuration, and with cardinal local edges but an oblique global configuration in **Fig. 1H**, and readers can judge for themselves which appears more dynamic. Readers can also compare diagonal **Fig. 1E** to vertical **Fig. 1G** with similar elements, and diagonal **Fig. 1H** to vertical **Fig. 1F** with similar elements, and judge for themselves which of each pair seems more dynamic.

Using stimuli that were systematic variations on this theme, we gained insight into the relative contributions of diagonal configuration versus oblique edges to perceived dynamism, providing evidence-based principles for eliciting dynamic percepts in visual art. In rating, ranking, and pupilometry experiments, we established the primacy of the orientation of the configuration over edges and identified ancillary structures that promote dynamism in still images.

## RESULTS

### Experiment 1: Rating dynamism of diagonals in still images

Experiment 1 quantitatively compared the role of local edges versus global configuration in creating a visual sense of dynamism. We generated a set of 60 geometric configurations, each consisting of a linear sequence of four identically shaped elements. The stimuli systematically varied in global configuration orientation, local edge orientation, element-size structure, and the direction of element-size variation. Twenty configurations contained equal-size elements (**Fig. 2A**), allowing us to isolate orientation-based cues, whereas the remaining forty had a wedge shape like Doesburg’s Composition with elements progressively increasing in size (**Fig. 2B**), adding a potential cue to depth or directional motion. In both tables, global configuration orientation varies across columns and local edge orientation varies across rows. 250 observers rated each stimulus on three perceptual dimensions—dynamism (static vs. dynamic), tridimensionality (flat vs. three-dimensional), and directionality (rising vs. falling) — using a −2, -1, 0, +1, +2 scale. Each cell displays the corresponding stimulus together with its mean dynamism rating averaged across participants; larger positive values indicate stronger perceived dynamism, whereas negative values indicate more static percepts.

**Fig. 2.**
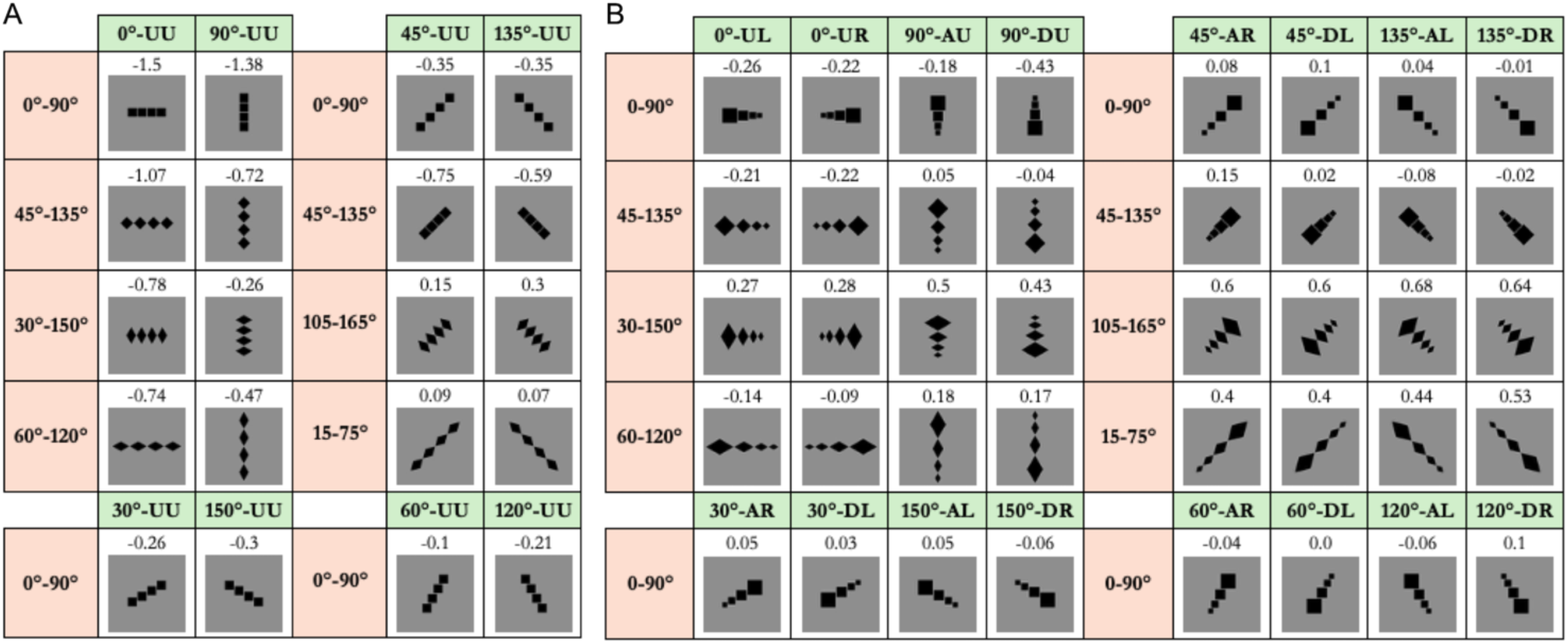
Experimental stimulus set. Stimuli are arranged according to global configuration orientation (columns) and local edge orientation (rows). A two-letter code specifies the pattern of element-size variation by combining the vertical component (A = Ascending, D = Descending, U = Uniform) with the horizontal component (R = Right, L = Left, U = Uniform). The numbers above each stimulus indicate the mean dynamism rating across the 250 observers in Experiment 1. **(A)** Configurations with equal-sized elements are designated as “UU”, i.e., uniform along both the horizontal and vertical axes. **(B)** Configurations with wedge-shaped elements, for which element size progressively increases along the direction specified by the size-pattern code (AR, AL, DR, DL, AU, DU, UR, UL).

**Fig. 3A** shows the number of responses for the five rating options (-2, -1, 0, +1, +2) for each stimulus and each scale. Stimuli are identified using the convention “Global orientation–Size pattern_Local edge orientations” by combining the codes in **Fig. 2** without the degree symbols, e.g. “0-AL_45-135” denotes a horizontal configuration (0°) with element size increasing upward and toward the left, and composed of elements with 45°–135° local edges. For the dynamism and tridimensionality scales, negative values indicate a perception of staticity or flatness, positive values indicate a stronger perception of dynamism or tridimensionality, and a score of 0 reflects uncertainty – that is, participants were unable to judge whether the image appeared more static or dynamic, or more flat or deep. In contrast, for the directionality scale, negative values indicate a predominantly rising impression and positive values indicate a predominantly falling impression, while a score of 0 means that the image is perceived as neither rising nor falling, but rather as stable.

**Fig. 3.**
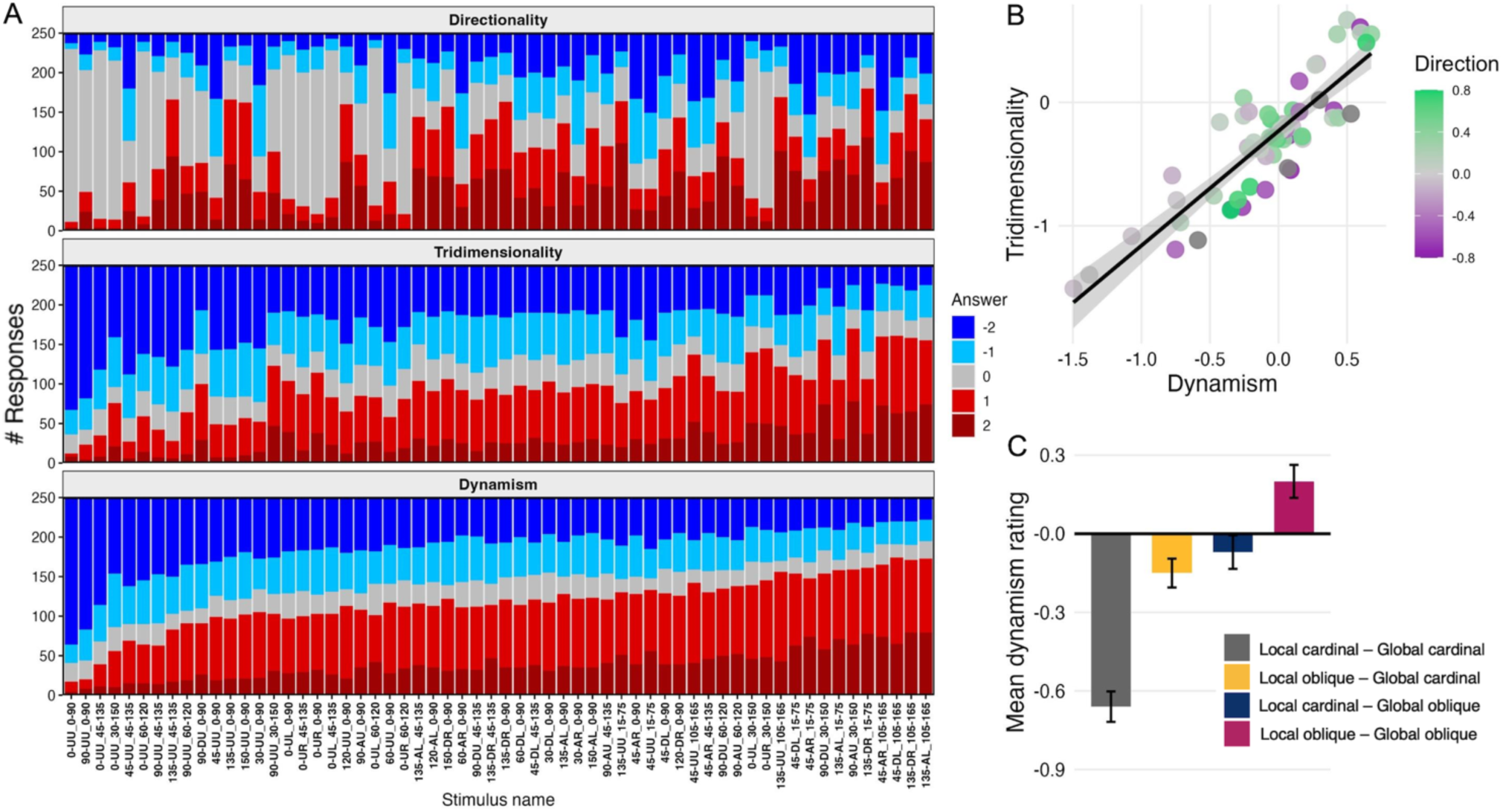
Relationships among perceptual dimensions and the contribution of local and global orientation to perceived dynamism. **(A)** Distribution of dynamism, three-dimensionality, and direction ratings across stimuli. Bars represent the number of responses in each rating category for every stimulus in Fig. 2, arranged according to increasing dynamism ratings (bottom panel). **(B)** Tridimensionality ratings versus Dynamism ratings. Points are colored according to Directionality ratings. The solid line shows the least-squares regression, with the shaded area representing the 95% confidence interval. **(C)**. Mean dynamism ratings for stimuli divided into four classes by global orientation crossed with local orientation. Error bars represent ± SEM across participants.

The dynamism scale reveals two stimuli with markedly lower scores compared to others. These are the two configurations with equal-size elements and with cardinal edges and cardinal global orientation: identical square elements arranged either vertically or horizontally (“0-UU_0-90” and “90-UU_0-90” in **Fig. 2A**). At the opposite end, four stimuli clearly show the highest dynamism, and all of them are wedge-shaped configurations with oblique edges and a diagonal configuration: diamond-shaped elements that increase in size either upward or downward along a ±45° diagonal axis (“135-AL_105-165”, “135-DR_105-165”, “45-DL_105-165”, “45-AR_105-165” in **Fig. 2B**).

For the tridimensionality scale, the stimuli eliciting the lowest and highest tridimensionality ratings are the same as those at the extremes of the dynamism scale, and the overall trend across stimuli is generally similar across the two dimensions. In both scales, stimuli with oblique orientations generally show positive response categories, suggesting that diagonal configurations evoke stronger dynamism and tridimensionality percepts. Only a small proportion of responses fall in the 0 category, indicating that participants rarely expressed uncertainty when judging how dynamic or how deep each image appeared.

The directionality scale showed a different pattern from the other two dimensions. Some wedge-shaped configurations with oblique orientations elicited predominantly rising or falling percepts, whereas configurations composed of equal-sized elements clustered around low-magnitude responses, with a marked prevalence of 0 ratings. The stronger directional impressions evoked by the wedge-shaped configurations may arise from the progressive size gradient, which can suggest depth or motion along the configuration axis, as well as from the asymmetry of the overall wedge shape. In contrast, equal-sized configurations provide few cues from which directional motion can be inferred, leading observers to judge them as neither rising nor falling. The absolute value of the directionality rating can therefore be interpreted as an index of perceived stabiliy/instability, with non-zero values indicating a tendency toward either rising or falling percepts and values close to zero indicating stable configurations. Consistent with this interpretation, the mean absolute directionality rating was substantially higher for oblique configurations (0.51) than for vertical (0.18) or horizontal (0.10) configurations.

Taken together, these observations suggest a close relationship between perceived dynamism, tridimensionality, and movement/instability. **Fig. 3B** summarizes the relationships among the three perceptual dimensions: each point corresponds to one image, plotted according to its mean dynamism and tridimensionality ratings, with colour indicating perceived directionality. The regression line highlights a strong linear relationship between dynamism and tridimensionality, and correlation analysis confirmed that images perceived as more dynamic were also judged as more three-dimensional (r = 0.87, 95% CI [0.79, 0.92], p < 0.001). In contrast, images with low dynamism and tridimensionality ratings are characterized by directionality values close to zero (grey symbols), whereas stimuli with higher dynamism and tridimensionality exhibit both positive (green) and negative (purple) directionality ratings. Accordingly, directionality did not correlate reliably with either dynamism (r = 0.10, p = 0.47) or tridimensionality (r = 0.06, p = 0.65). The absolute value of the directionality ratings, interpreted as an index of perceived stability, showed a modest but significant positive correlation with dynamism (r = 0.26, 95% CI [0.004, 0.48], p = 0.047), but was unrelated to tridimensionality (r = –0.03, p = 0.83), indicating that this measure dissociates perceived instability from perceived depth.

Having clarified how the three perceptual dimensions relate to one another, we next focus on the critical dynamism dimension. **Fig. 3C** shows the mean dynamism ratings for the four stimulus categories that combine cardinal or oblique local edge orientation with cardinal or oblique global configurational orientation. The two global oblique categories were rated as noticeably more dynamic (−0.07 ± 0.064 and 0.20 ± 0.063) than the two global cardinals (-0.66 ± 0.058 and -0.15 ± 0.055). Oblique edge orientations exert a similar but smaller effect, enhancing the dynamism of both classes of global configurations.

Although the pattern shown in **Fig. 3C** is convincing, the stimulus categories aggregate images that differ along multiple visual dimensions. To isolate the contribution of individual features – global orientation, local edge orientation, element spacing, and direction of element size variation – we performed a series of planned contrasts between the perceived dynamism ratings of targeted stimuli. The complete statistical results are reported in ***SI Appendix***, **Table S1**, and detailed descriptions of each comparison are provided in the supplementary text. The main findings are summarized below.

The contribution of global configuration orientation emerged as the strongest determinant of perceived dynamism: diagonal configurations were consistently rated as more dynamic than cardinal configurations, regardless of whether local edges were oblique or cardinal (both *p* < .05). Among cardinal configurations, vertical layouts were generally perceived as slightly more dynamic than horizontal ones, whereas no reliable differences emerged between the two mirror-symmetric oblique orientations (45° vs. 135°), indicating isotropy within the oblique axis.

The comparisons designed to assess local edge orientation effects revealed a more complex pattern. Within cardinal configurations, stimuli with oblique local edges were judged as more dynamic than those with cardinal edges, whereas within oblique configurations the opposite pattern emerged, with cardinal-edged stimuli rated as more dynamic than oblique-edged stimuli (both *p* < .01). The edge effects were complicated by configurations with the more widely separated elements receiving higher dynamism ratings: oblique-edged diamonds in cardinal configurations and cardinal-edged squares in oblique configurations, suggesting an effect of spatial dispersion, a more general result than in **Fig. 3C**, based on more classes of stimuli. To test this possibility directly, we compared more widely dispersed and more compact configurations, confirming that inter-element spacing itself enhances perceived dynamism (*p* < .001). The same effect generalized across different element shapes, indicating that increased spacing contributes to perceived dynamism independently of the specific geometry of the elements.

Having established that inter-element spacing contributes independently to perceived dynamism, we next compared configurations matched for spacing to isolate the relative contributions of global and local orientation. Globally oblique configurations with cardinal local edges were rated as more dynamic than globally cardinal configurations with oblique local edges (*p* < .001), confirming that global configuration orientation contributes more strongly than local edge orientation. Finally, the difference was even greater when both global and local orientations were oblique than when both were cardinal (*p* < .001), indicating that oblique local edges provide a smaller but significant additional contribution to perceived dynamism.

Another result we did not foresee was the effect of element-size variation. Wedge-shaped configurations were consistently rated as more dynamic than otherwise similar equal-size configurations across all global orientations (all *p* < .001). In contrast, the direction of size progression itself (e.g., leftward vs. rightward, ascending vs. descending) produced little or no effect, except for vertical configurations, in which top-heavy arrangements were judged slightly more dynamic than bottom-heavy ones.

Overall, these planned comparisons show that perceived dynamism is determined primarily by configuration orientation, is enhanced by greater inter-element spacing and systematic size gradients that create wedge shapes, and is only secondarily influenced by local edge orientation.

### Experiment 2: Paired comparison rankings and a perceptual scale for perceived dynamism

Experiment 1 asked observers to assign absolute ratings to individual stimuli and provided a robust identification of the physically static visual cues contributing to the perception of dynamism. However, absolute ratings are often more strongly influenced by individual response criteria than relative rankings (13), and because Experiment 1 was conducted online, viewing conditions could not be standardized across participants. Experiment 2 therefore complemented these findings by measuring perceived dynamism through pairwise comparisons under controlled laboratory conditions, and allowed us to derive a perceptual scale of dynamism (14).

From the stimulus set shown in **Fig. 2**, we created 168 stimulus pairs designed to isolate four visual factors hypothesized to contribute to perceived dynamism: (i) global configuration orientation (***SI Appendix,* Table S2**), (ii) local edge orientation (***SI Appendix,* Table S3**), (iii) direction of element-size variation (***SI Appendix,* Table S4**), and (iv) the interaction between local and global orientation (***SI Appendix,* Table S5**). Within each subset, one visual factor was manipulated while the remaining factors were held constant.

On each trial, two images were presented side by side simultaneously, allowing observers to directly compare their perceived dynamism. 50 observers ranked each pair using a 5-point scale ranging from –2 (the stimulus on the left appeared more dynamic) to +2 (the stimulus on the right appeared more dynamic), with 0 indicating equal perceived dynamism. Each pair was presented six times, with the left–right positions of the two stimuli counterbalanced across repetitions.

To assess the consistency between the two measurement methods, we correlated the difference in mean dynamism ratings from Experiment 1 (***SI Appendix***, **Table S6**) with the corresponding mean pairwise dynamism ranking from Experiment 2 (***SI Appendix***, **Table S7**). Pairs were grouped according to the specific visual factor they were designed to probe — global configuration orientation, local edge orientation, the direction of element size variation, and the interaction between global and local orientations — and correlations were computed separately for each subset (**Fig. 4A**). This analysis revealed strong and significant monotonic relationships for pairs differing in global configuration orientation (ρ = 0.804, p < .001), local edge orientation (ρ = 0.787, p < 0.001), and the interaction between global and local orientation (ρ = 0.709, p = 0.01). In contrast, there was little spread of dynamism rating difference or pairwise score for direction of element size variation, so the correlation was not significantly different from zero (ρ = -0.089, p = 0.708). Thus, stimulus manipulations that strongly modulated perceived dynamism in the rating-based task produced consistent biases in relative judgments. After showing that orientation-based effects on perceived dynamism are robust across measurement paradigms, we used pairwise comparisons to create a perceptual scale for perceived dynamism as a function of configuration and edge orientations.

**Fig. 4.**
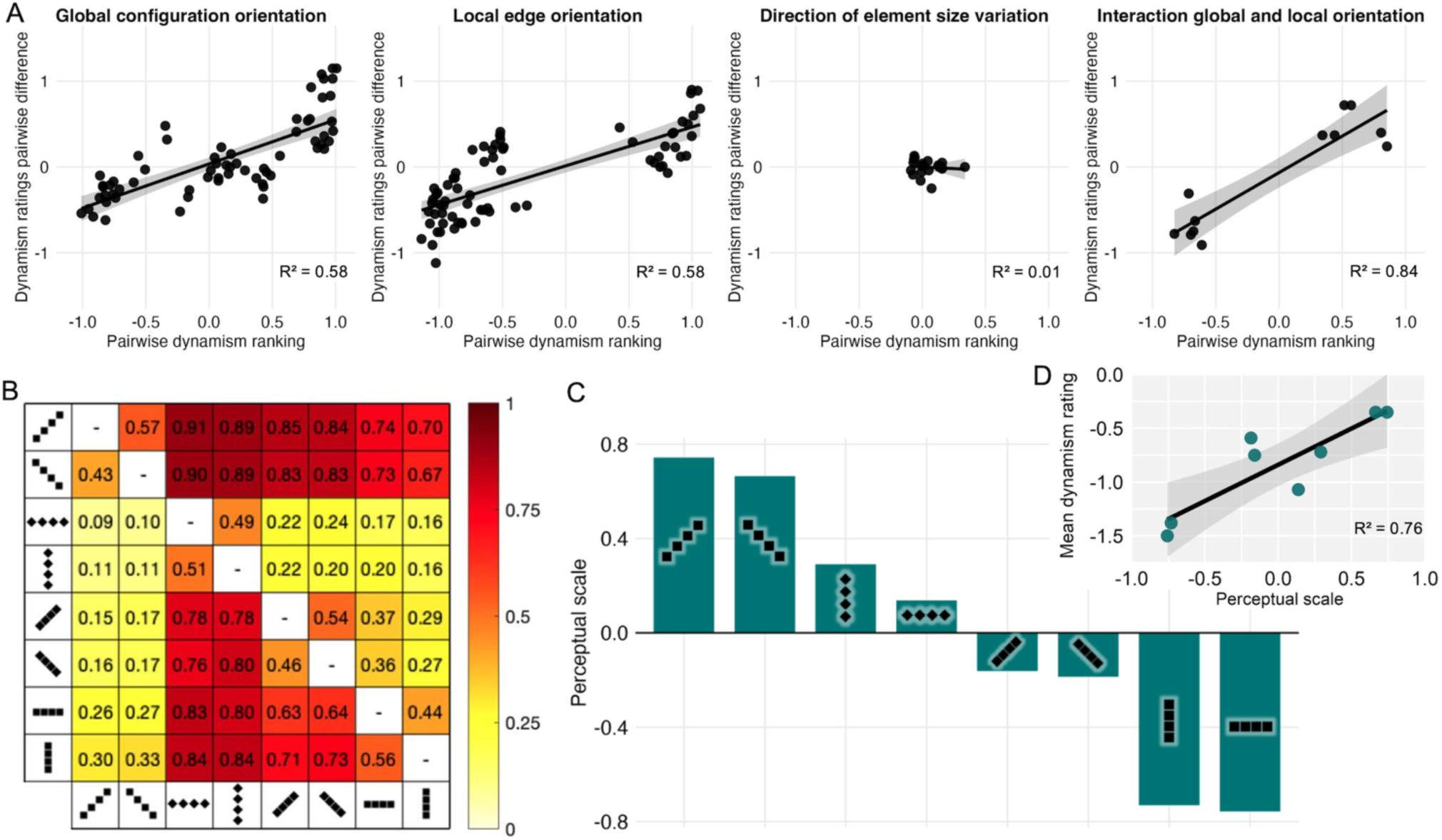
Pairwise comparisons and a perceptual scale of dynamism. **(A)** For each pair of stimuli, the relationship of the mean difference between dynamism ratings (Experiment 1) to pairwise dynamism judgments (Experiment 2). Stimulus pairs are grouped according to the visual factor manipulated. The solid lines show least-squares linear fits. The coefficient of determination (R²) from the linear fit is reported in the bottom-right corner of each panel. **(B)** Proportion of times the row stimulus was ranked as more dynamic than the column stimuli. **(C)** Eight stimuli placed on the derived perceptual scale. **(D)** Correlation between perceptual scale and dynamism ratings in Experiment 1.

We focused on a subset of eight stimuli for which all possible pairwise combinations were tested (28 pairs in total); **Fig. 4B** shows the comparison matrix derived from participants’ responses. Each row indicates the probability that the row stimulus was perceived as more dynamic than each of the column stimuli, whereas each column reflects the probability that the column stimulus was judged as less dynamic than each row stimulus.

Using this matrix, we applied an extended maximum-likelihood implementation of Thurstone’s scaling method (14) to derive a perceptual scale of perceived dynamism (**Fig. 4C**). In this scale, each stimulus is assigned a position along a latent psychological dimension reflecting its perceived dynamism. Higher values indicate more dynamic stimuli, and distances between scale values reflect the relative perceptual differences in dynamism between stimuli. Interestingly, the perceptual scale also revealed a clear effect of inter-element spacing, with configurations containing larger spacing consistently positioned higher on the scale. This effect had not been considered during stimulus design and independently replicated the pattern observed in Experiment 1. Within similar spacings, the oblique configurations are more dynamic than the cardinals. As shown in **Fig. 4D**, the perceptual scale strongly correlated with the mean dynamism ratings obtained in Experiment 1 (ρ = 0.83, p < .05).

To the extent that the perceptual scale is a good fit to the paired-comparisons, it indicates that the scale measures exactly one uni-dimensional latent trait. Perceived dynamism in static images can thus be treated as a linear psychological continuum, with distances on the scale representing subjective differences in a possible internal representation. Due to limitations on the number of pairs that could be feasibly compared with multiple repeats, the scale was derived for just the 8 configurations in **Fig. 4C**, but there is no reason to rule out that the other configurations would also fit on this scale, probably close to positions predicted by the dynamism ratings, given the strong correlation in **Fig. 4D**.

### Experiment 3: Pupil responses to perceived dynamism

Ratings and rankings involve conscious judgments. To see whether diagonals also evoke involuntary signatures of neural responses that are different from responses to cardinals, we measured pupil responses to 13 different configurations that gave the highest and lowest perceived dynamism ratings in Experiment 1 (**Fig. 5B** Legend). Although pupil responses are primarily driven by luminance, they are also modulated by a wide range of perceptual, cognitive, and emotional factors (15–21). Of particular interest are findings showing that the pupil dilates in response to moving objects (22), suggesting that pupillary responses are sensitive to motion-related visual processing and may reflect increased arousal associated with dynamic visual events (23). Relevant to the present study, pupil size has also been shown to be modulated by the perception of implied and illusory motion in static images (22, 24, 25).

**Fig. 5.**
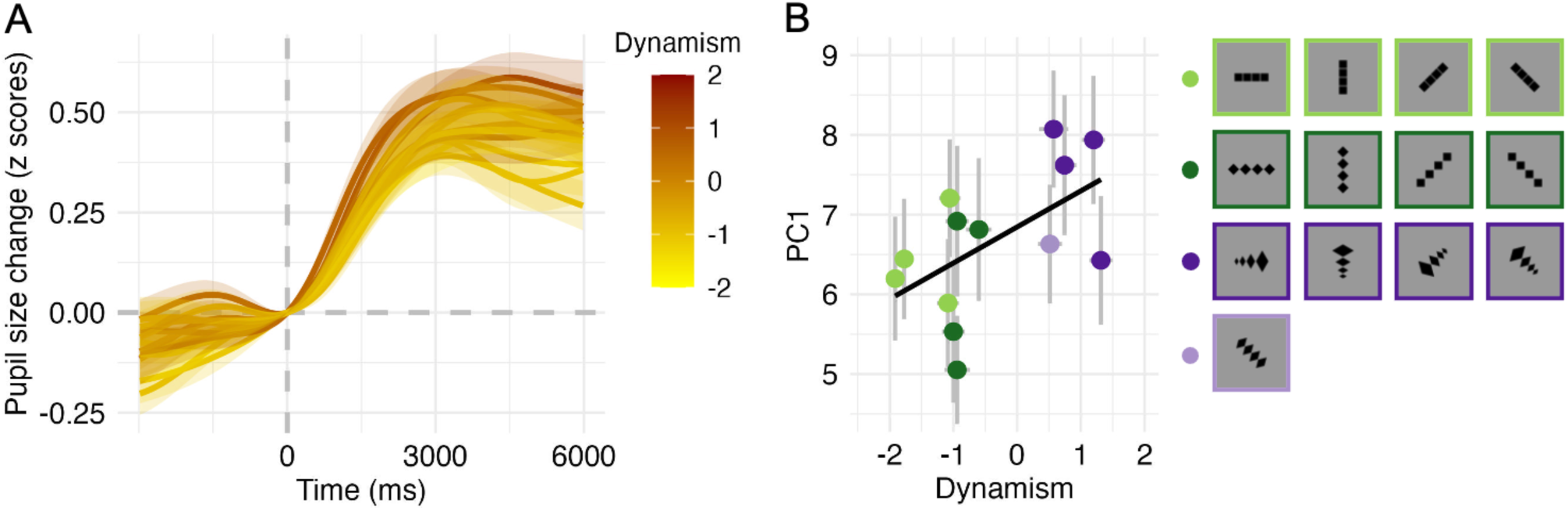
Pupil responses as a function of perceived dynamism. **(A)** Time course of the pupillary response for all stimuli. The color of each curve and its corresponding shaded error band reflects the mean perceived dynamism rating obtained during the final viewing session, with warmer colors indicating higher perceived dynamism. **(B)** Relationship between perceived dynamism and the first principal component (PC1) of the pupillary response. Error bars represent ± SEM across participants. The legend shows the 13 stimuli used in this experiment binned by category.

Stimuli (7°x7°) were presented centrally for 6 s while pupil size was recorded. From the static rated stimuli, we used 4 configurations with congruent local and global orientations and 4 with incongruent orientations. From the dynamic rated configurations, we used 4 wedge-shapes spanning the same configuration orientations as the static set, and one with equal-size elements. The inter-stimulus interval varied between 3 and 4 s randomly and was filled with a scrambled baseline stimulus composed of unstructured patterns that contained approximately the same number of black pixels as the experimental stimuli. Participants were instructed to maintain central fixation throughout stimulus presentation. Each stimulus was repeated eight times. In an additional session, each stimulus was presented once more at a larger size (21°x21°), and participants were allowed to freely explore the images for 8 s and asked to rate each configuration along the dynamism, tridimensionality, and direction dimensions of Experiment 1, using the same 5-point scale. The ratings for each subject could be directly related to their pupillary responses. Mean dynamism ratings across stimuli showed a strong correlation with mean ratings from Experiment 1 (ρ = 0.94, p < .001).

**Fig. 5A** shows the time course of the pupillary response for all stimuli, with color saturation reflecting the average dynamism ratings. Pupil size exhibited a gradual increase over time for all conditions. This sustained dilation is likely driven by multiple factors. The scrambled baseline stimulus preceding each image was designed to match the target stimuli in mean luminance to reduce the chances of pupil dilation due to a luminance decrement, but we cannot rule out some contribution to the observed pupil enlargement by the spatially organized, high-contrast dark elements present in the stimuli. In addition, the gradual increase in pupil size may reflect ongoing visual and cognitive processing elicited by viewing the stimuli, rather than solely a transient pupillary light reflex. However, since all configurations were matched in overall area and average luminance, any differences in pupil responses across stimuli are not due to differences in luminance.

Particularly during the later phases of the pupil response, configurations rated as more dynamic (indicated by the color gradient) tended to elicit stronger dilation. To quantify these differences, we applied principal component analysis (PCA) to the pupil time-course (26). The first principal component (PC1) accounted for 82% of the total variance and captured the dominant dilation pattern elicited by the stimuli. PC1 scores were therefore used as a summary measure of pupillary response magnitude. A repeated-measures ANOVA revealed a significant effect of configuration on PC1 scores (F(12,408) = 1.83, p = 0.042), indicating that the magnitude of the pupillary response differed across the 13 configurations.

We tested whether this variation was related to perceived dynamism by computing, for each participant, the Spearman correlation between dynamism ratings of each configuration and the corresponding PC1 scores (average ρ = 0.13, SEM = 0.04). A one-sample t-test showed that these correlation coefficients were significantly greater than zero (t(34) = 2.69, p = 0.011), indicating that stronger pupillary responses were consistently associated with higher perceived dynamism across participants. **Fig. 5B** illustrates this relationship at the stimulus level: each point represents one configuration and colors indicate the stimulus category. The black least-squares linear regression line shows a positive but modest correlation (r = 0.56), identifying pupil size as an objective physiological marker of dynamism perception.

Taken together, the three experiments demonstrate that perceived dynamism constitutes a coherent internal dimension, rooted in specific geometric properties of static images and accompanied by a measurable psychophysiological response.

## DISCUSSION

Representing movement in static art has been a challenge since the beginning of cave drawings (27). Arnheim pointed out that the image of a waterfall may not look particularly dynamic, so perceived dynamism does not always derive from associating an object with its intrinsic motion from memory, but by features such as asymmetry with an indication of direction (28). Diagonals and wedges have been claimed to suggest motion, as exemplified by El Lissitzky’s Beat the Whites with the Red Wedge (1919). Our three experiments confirm empirically that human observers can perceive different levels of dynamism in still images which are quite simple and devoid of semantic content.

The first rating experiment showed that dynamism is conveyed by diagonal configurations, and that dynamism is enhanced by wedge-shaped configurations, but more by elements separated at vertices forming loose wedges than by solid-appearing wedges formed by elements separated at edges. The primacy of diagonals was corroborated by the second paired ranking experiment, which did not rely on observers assigning numerical values to their evaluations but ranking which image of a pair of images was higher in dynamism. The design of this experiment provided stronger evidence but was limited to testing fewer factors. Dynamism goes beyond being solely a perceptual quality to also involving cognition and possibly even emotion. Human pupil dilation has been shown to be an involuntary correlate of perceptual, cognitive, and emotional factors (15–21), potentially providing a measure of perceived dynamism that is not subject to artifacts in rating and ranking methods, and the third experiment showed a modest but positive correlation between pupil dilation and perceived dynamism, consistent with previous results correlating individual reports of dynamism across real, illusory, and implied motion with individual pupil responses (22, 24).

Given the main results of this study, it is worth revisiting the four ideas we considered in the Introduction as possible explanations for the power of diagonals despite the oblique effect. Two of the ideas concerned configurations, not edges, and we provide some evidence for their validity. It has been conjectured that diagonal figures could generate percepts of depth or tridimensionality, allowing figures to appear dynamic as if they were emerging out of the picture plane. Experiment 1 results support this conjecture as the correlation between dynamism and tridimensionality ratings was 0.87. It has also been noted that real-world objects that are neither vertical nor horizontal are often free-falling or moving intentionally, which could associate an intrinsic dynamic meaning with diagonal configurations. The absolute value of the directionality rating gives a measure of imagined movement irrespective of upwards or downwards, and its higher value averaged for all diagonal configurations was substantially higher than for vertical and horizontal configurations, providing support to the idea of movement being associated with diagonal configurations. There has been little investigation of how global orientations and shapes of multi-element configurations are extracted, and how that contributes to sensitivity, salience or surprise. This study suggests that such investigations are needed to understand scene perception and its cognitive and emotive components.

One intriguing aspect of the results is that although the contribution of oblique local edges, which would be subject to the oblique effect, is smaller than that of global configurations, they also contribute to percepts of dynamism. The oblique effect has been demonstrated against spatially uniform backgrounds, but for more complex situations we conjectured in the Introduction that novelty detection could be enhanced by adaptation to stimulus spatial statistics, making the more frequent cardinal orientations less salient and potentially causing oblique edges or contours to stand out, and that when most local orientations in an image are cardinal, the implicit frame they form could further highlight oblique features. These conjectures are compatible with results showing greater matched salience for obliquely oriented components of broadband stimuli with naturalistic spatial statistics (29). The salience-enhancing mechanism proposed by this study was divisive surround normalization in primary visual cortex, but this is incompatible with multiple aspects of orientation processing in V1 (30), so adaptation and flashed salience experiments may help identify the neural mechanisms.

Taken together, these considerations suggest in multiple ways that associating diagonals with dynamism may have more to do with extracted qualities such as salience than with the limitations associated with encoding oblique orientations in early cortical processing. That we find a coherent perceptual scale for dynamism that correlates well with ratings suggests the existence of internal representations of dynamism. In addition, finding systematic pupil dilations as a function of perceived dynamism reinforces the idea that diagonals generate salience, surprise, or related qualities that have been shown to influence pupil responses.

Visual artists often work out perceptual puzzles that can inspire scientists studying visual perception; for example, Salvador Dalí’s Agnostic Symbol (1932) uses a detached shadow to make a spoon appear to hover over the desert, presaging demos of objects appearing to change elevation by moving apart from their shadows (31). There are also many examples in the other direction; for example, Musatti’s stereokinetic experiments with rotating disks (32) were used by Duchamp in his Rotoreliefs. At a more abstract level, the use of diagonals to create dynamism by artists and photographers inspired us to think past the oblique effect and to consider processes that go beyond mechanisms that efficiently encode environmental statistics. In return, we offer artists an empirical grounding for micro-strategies for manipulating dynamism by using diagonal and wedge-shaped configurations with visually separated elements. This study is the first step in understanding how the visual system extracts properties of extended multi-element spatial configurations and uses them for salience, drama, and emotional responses.

## MATERIALS AND METHODS

More details of the experimental procedures, stimuli, and statistical analyses are provided in the **SI Appendix**.

This study was approved by the local ethics committee (“Commissione per l’Etica della Ricerca”, University of Florence, January 17, 2024, No. 292) and conducted in accordance with the Declaration of Helsinki. Participants in all three experiments were naïve to the purpose of the study, had normal or corrected-to-normal vision, and provided informed consent before participation.

Main statistical analyses were performed in R (version 4.4.2) and MATLAB (R2023b).

### Experiment 1

250 adults (mean age = 25.4 years, SD = 9.6) rated the 60 stimuli shown in **Fig. 2**. The study was conducted online using Google Forms. On each page, a single stimulus was displayed, and participants rated perceived dynamism, tridimensionality, and direction using three 5-point scales (−2 to +2). Equal-size stimuli were presented first, followed by wedge-shaped stimuli, with randomized stimulus order within each block. The order of the rating scales was randomized across participants.

Responses were recoded so that higher values consistently reflected greater perceived dynamism (and analogously for the other scales). Stimulus ratings were averaged across participants. Because ratings were ordinal and deviated from normality, statistical significance was assessed using the Wilcoxon tests, whereas t-statistics are reported as descriptive measures of effect size and direction. Differences in the distributions of categorical responses were additionally assessed using chi-square tests.

### Experiment 2

From the original 60 stimuli, we generated 168 stimulus pairs designed to isolate the effects of global configuration orientation, local edge orientation, element-size variation, and their interactions (**SI Appendix, Tables S2–S5**). 50 adults (mean age = 25.2 years, SD = 6.9) made pairwise comparisons in a controlled lab setting. On each trial, two stimuli were presented simultaneously until response, and participants indicated which configuration appeared more dynamic using a 5-point comparative scale. Each stimulus pair was presented six times with counterbalanced left-right positions, yielding 1008 trials presented in randomized order across six blocks.

### Experiment 3

35 adults (mean age = 23.1 years, SD = 2.4) participated in a pupillometry experiment. Based on the ratings in Experiment 1, we selected a subset of 13 representative stimuli spanning the range from static to dynamic configurations (Fig. 5B). The experiment comprised two phases. During the first phase, participants maintained central fixation while the 13 stimuli were each presented eight times following a scrambled baseline stimulus, allowing stimulus-evoked pupil responses to be recorded. During the second phase, participants freely viewed each stimulus once before rating its perceived dynamism, tridimensionality, and direction using the same scales as in Experiment 1. Eye position and pupil size were recorded continuously with an EyeLink 1000 eye tracker at 500 Hz. Pupil recordings were preprocessed to remove blinks and eye movements, baseline-corrected, normalized within participants, and summarized using principal component analysis (PCA).

## ACKNOWLEDGMENTS

QZ was supported by NEI grants EY035085 and EY035838. SC and EB were partially supported by the co-funding of the European Union - Next Generation EU, in the context of The National Recovery and Resilience Plan, Mission 4, Component 2, Investment 1.5 Ecosystems of Innovation, Project Tuscany Health Ecosystem (THE), ECS00000017, Spoke 3, CUP: B83C22003920001. EB was also partially supported by the European Union - NextGenerationEU, Investment line 1.2 “Funding projects presented by young researchers” (BEST-VS), CUP: E73C25000210001.

## DATA, MATERIALS, AND SOFTWARE AVAILABILITY

Data, experimental materials, and analysis code supporting the findings of this study are publicly available on Zenodo at https://doi.org/10.5281/zenodo.21773469.

## Notes

https://doi.org/10.5281/zenodo.21773469

### Competing Interest Statement

The authors have declared no competing interest.

### Summary of Updates

Updated the data availability statement and made minor revisions to the final paragraph of the Discussion.

