## Supplementary Material for "Dynamism in paintings and photographs: how the power of diagonals in art overcomes the oblique effect in visual perception"

**Corresponding author:** Qasim Zaidi

**This PDF file includes:**

Tables S1 to S7

Supporting text for Table S1

Supporting text for Methods and Materials

SI References

**Table S1. Experiment 1: statistical comparisons between stimulus groups.** Results of the planned comparisons performed on six stimulus sets designed to isolate the contribution of specific visual features to perceived dynamism, including global configuration orientation, local edge orientation, element size variation, and their interactions. For each stimulus set, the left column illustrates the configurations included in the comparison, whereas the table reports the contrasted stimulus groups together with the corresponding t-statistic from the paired-samples t-test, the p-value from the Wilcoxon signed-rank test (used to assess statistical significance), and the $\chi^2$ statistic with its associated p-value from the chi-square test (see *Materials and Methods* for details). Detailed descriptions and interpretations of the individual comparisons are provided in the Supplementary Results below the table.

| | N | Group 1 | Group 2 | *t*-value, *p*-Wilcoxon | $X^2$-value, *p*-value |
|---|---|---|---|---|---|
| **Set 1** 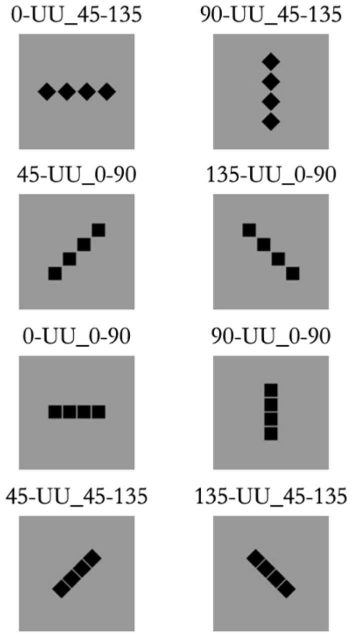  | 1.1 | 0-UU_45-135 & 90-UU_45-135 | 45-UU_45-135 & 135-UU_45-135 | -2.89, 0.01 | 14.68, 0.005 |
| | 1.2 | 0-UU_0-90 & 90-UU_0-90 | 45-UU_0-90 & 135 UU_0-90 | -13.9, <.001 | 194.28, <.001 |
| | 1.3 | 0-UU_45-135 & 90-UU_45-135 | 0-UU_0-90 & 90-UU_0-90 | 9.08, <.001 | 68.87, <.001 |
| | 1.4 | 45-UU_0-90 & 135-UU_0-90 | 45-UU_45-135 & 135-UU_45-135 | 4.39, <.001 | 15.04, 0.005 |
| | 1.5 | 0-UU_45-135 & 90-UU_45-135 | 45-UU_0-90 & 135-UU_0-90 | -7.41, <.001 | 46.43, <.001 |
| | 1.6 | 45-UU_45-135 & 135-UU_45-135 | 0-UU_0-90 & 90-UU_0-90 | 10.18, <.001 | 114.01, <.001 |
| | 1.7 | 0-UU_45-135 & 90-UU_45-135 & 45-UU_0-90 & 135-UU_0-90 | 0-UU_0-90 & 90-UU_0-90 & 45-UU_45-135 & 135-UU_45-135 | 8.66, <.001 | 65.27, <.001 |
| | 1.8 | 0-UU_45-135 | 90-UU_45-135 | -4.18, <.001 | 15, 0.005 |
| | 1.9 | 0-UU_0-90 | 0-UU_0-90 | -1.8, 0.068 | 6.53, 0.163 |
| | 1.10 | 45-UU_0-90 | 135-UU_0-90 | 0, 0.984 | 2.22, 0.695 |
| | 1.11 | 45-UU_45-135 | 135-UU_45-135 | -1.95, 0.067 | 2.04, 0.729 |
| **Set 2** 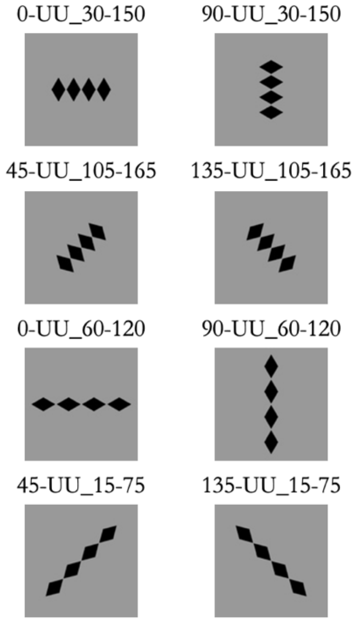  | 2.1 | 0-UU_30-150 & 90-UU_30-150 | 45-UU_105-165 & 135-UU_105-165 | -10.66, <.001 | 78.52, <.001 |
| | 2.2 | 0-UU_60-120 & 90-UU_60-120 | 45-UU_15-75 & 135-UU_15-75 | -8.93, <.001 | 62.55, <.001 |
| | 2.3 | 0-UU_30-150 | 90-UU_30-150 | -5.64, <.001 | 23.51, <.001 |
| | 2.4 | 0-UU_60-120 | 90-UU_60-120 | -2.71, 0.007 | 9.41, 0.052 |
| | 2.5 | 0-UU_30-150 & 90-UU_30-150 & 45-UU_105-165 & 135-UU_105-165 | 0-UU_60-120 & 90-UU_60-120 & 45-UU_15-75 & 135-UU_15-75 | 2.43, 0.028 | 10.51, 0.033 |

<table>
<tr><td>Set 3<br>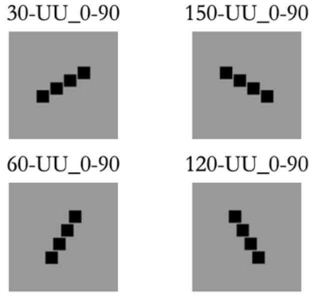<br></td><td>3.1</td><td>30-UU_0-90 &<br>150-UU_0-90</td><td>60-UU_0-90 &<br>120-UU_0-90</td><td>-2.23,<br>0.032</td><td>2.37,<br>0.667</td></tr>
</table>

<table>
<tr><td rowspan="2">Set 4<br>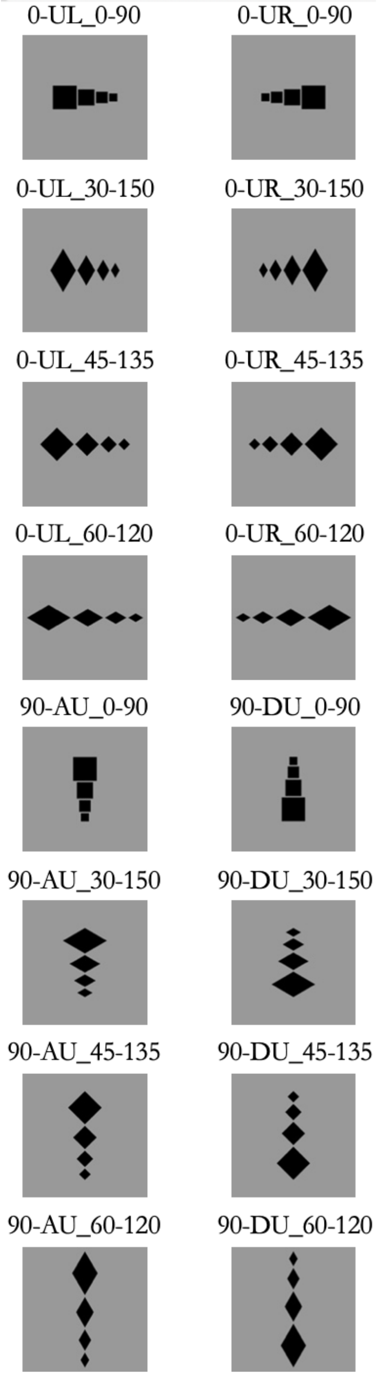<br></td><td>4.1</td><td>0-UL_0-90 &<br>0-UL_30-150 &<br>0-UL_45-135 &<br>0-UL_60-120</td><td>0-UR_0-90 &<br>0-UR_30-150 &<br>0-UR_45-135 &<br>0-UR_60-120</td><td>-0.54,<br>0.589</td><td>1.98,<br>0.74</td></tr>
<tr><td>4.2</td><td>90-AU_0-90 &<br>90-AU_30-150 &<br>90-AU_45-135 &<br>90-AU_60-120</td><td>90-DU_0-90 &<br>90-DU_30-150 &<br>90-DU_45-135 &<br>90-DU_60-120</td><td>2.78,<br>0.003</td><td>3.19,<br>0.526</td></tr>
</table>

| | | | | | |
|---|---|---|---|---|---|
| **Set 5**<br>30-AR_0-90 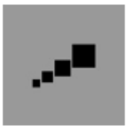 30-DL_0-90 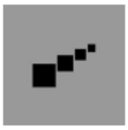<br>45-AR_0-90 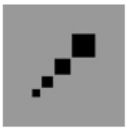 45-DL_0-90 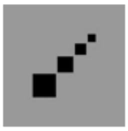<br>45-AR_15-75 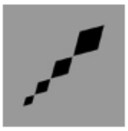 45-DL_15-75 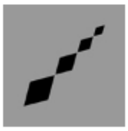<br>45-AR_45-135 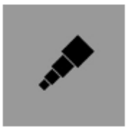 45-DL_45-135 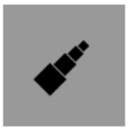<br>45-AR_105-165 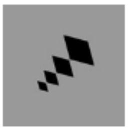 45-DL_105-165 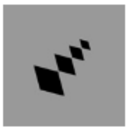<br>60-AR_0-90 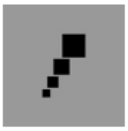 60-DL_0-90 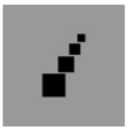<br>150-AL_0-90 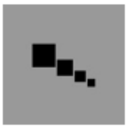 150-DR_0-90 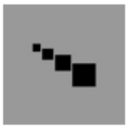<br>135-AL_0-90 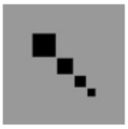 135-DR_0-90 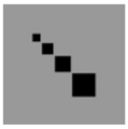<br>135-AL_15-75 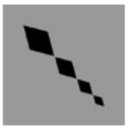 135-DR_15-75 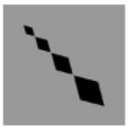<br>135-AL_45-135 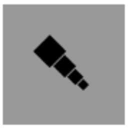 135-DR_45-135 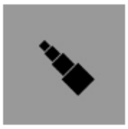<br>135-AL_105-165 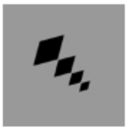 135-DR_105-165 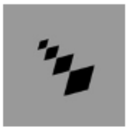<br>120-AL_0-90 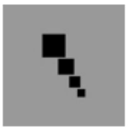 120-DR_0-90 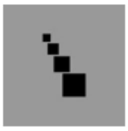 | 5.1 | 30-AR_0-90 &<br>45-AR_0-90 &<br>45-AR_15-75 &<br>45-AR_45-135 &<br>45-AR_105-165 &<br>60-AR_0-90 &<br>150-AL_0-90 &<br>135-AL_0-90 &<br>135-AL_15-75 &<br>135-AL_45-135 &<br>135-AL_105-165 &<br>120-AL_0-90 | 150-DR_0-90 &<br>135-DR_0-90 &<br>135-DR_15-75 &<br>135-DR_45-135 &<br>135-DR_105-165 &<br>120-DR_0-90 &<br>30-DL_0-90 &<br>45-DL_0-90 &<br>45-DL_15-75 &<br>45-DL_45-135 &<br>45-DL_105-165 &<br>60-DL_0-90 | -0.12,<br>0.621 | 2.64,<br>0.619 |

| | | | | | |
|---|---|---|---|---|---|
| **Set 6**<br>0-UU_0-90 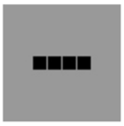 0-UL_0-90 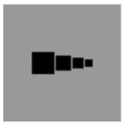 0-UR_0-90 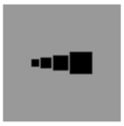<br>0-UU_45-135 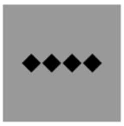 0-UL_45-135 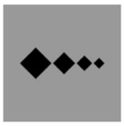 0-UR_45-135 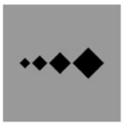<br>0-UU_30-150 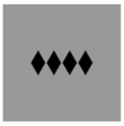 0-UL_30-150 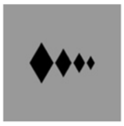 0-UR_30-150 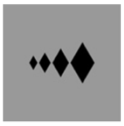<br>0-UU_60-120 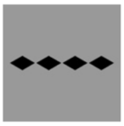 0-UL_60-120 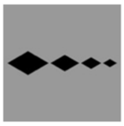 0-UR_60-120 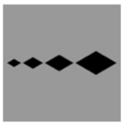 | 6.1 | 0-UU_0-90 &<br>0-UU_45-135 &<br>0-UU_30-150 &<br>0-UU_60-120 | 0-UL_0-90 &<br>0-UL_45-135 &<br>0-UL_30-150 &<br>0-UL_60-120 &<br>0-UR_0-90 &<br>0-UR_45-135 &<br>0-UR_30-150 &<br>0-UR_60-120 | -15.21,<br><.001 | 307.89,<br><.001 |
| 90-UU_0-90 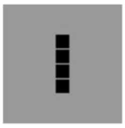 90-AU_0-90 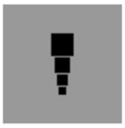 90-DU_0-90 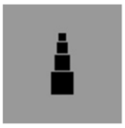<br>90-UU_45-135 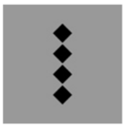 90-AU_45-135 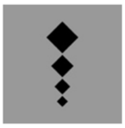 90-DU_45-135 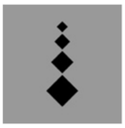<br>90-UU_45-135 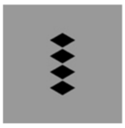 90-AU_30-150 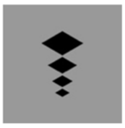 90-DU_30-150 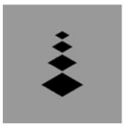<br>90-UU_60-120 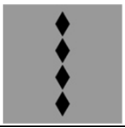 90-AU_60-120 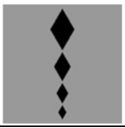 90-DU_60-120 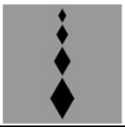 | 6.2 | 90-UU_0-90 &<br>90-UU_45-135 &<br>90-UU_30-150 &<br>90-UU_60-120 | 90-AU_0-90 &<br>90-AU_45-135 &<br>90-AU_30-150 &<br>90-AU_60-120 &<br>90-DU_0-90 &<br>90-DU_45-135 &<br>90-DU _30-150 &<br>90-DU _60-120 | -13.08,<br><.001 | 197.89,<br><.001 |

| | | | | | |
|---|---|---|---|---|---|
| 45_45-135_UU, 45_45-135_AR, 45_45-135_DL<br>45_0-90_UU, 45_0-90_AR, 45_0-90_DL<br>45_105-165_UU, 45_105-165_AR, 45_105-165_DL<br>45_15-75_UU, 45_15-75_AR, 45_15-75_DL<br>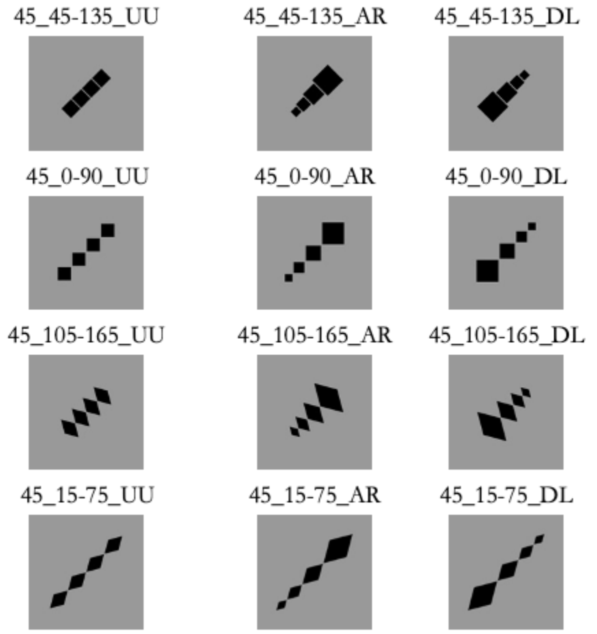 | 6.3 | 45-UU_45-135 & 45-UU_0-90 & 45-UU_105-165 & 45-UU_15-75 | 45-AR_45-135 & 45-AR_0-90 & 45-AR_105-165 & 45-AR_15-75 & 45-DL_45-135 & 45-DL_0-90 & 45-DL_105-165 & 45-DL_15-75 | -8.65, <.001 | 92.61, <.001 |
| 135_45-135_UU, 135_45-135_AL, 135_45-135_DR<br>135_0-90_UU, 135_0-90_AL, 135_0-90_DR<br>135_105-165_UU, 135_105-165_AL, 135_105-165_DR<br>135_15-75_UU, 135_15-75_AL, 135_15-75_DR<br>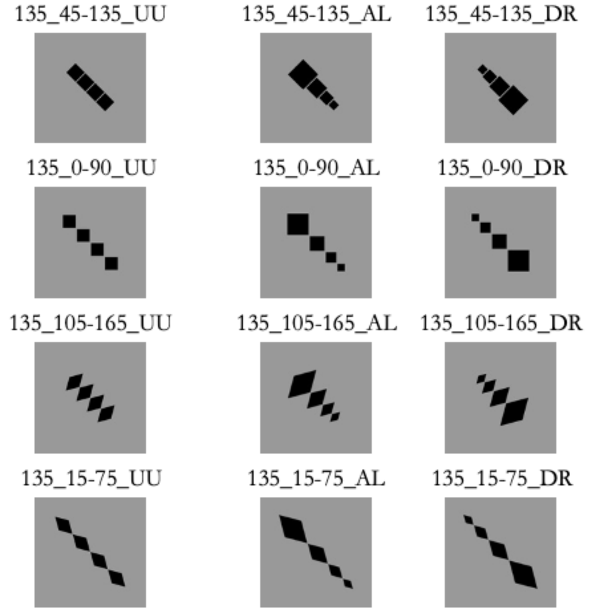 | 6.4 | 135-UU_45-135 & 135-UU_0-90 & 135-UU_105-165 & 135-UU_15-75 | 135-AL_45-135 & 135-AL_0-90 & 135-AL_105-165 & 135-AL_15-75 & 135-DR_45-135 & 135-DR_0-90 & 135-DR_105-165 & 135-DR_15-75 | -6.8, <.001 | 64.88, <.001 |

### Supporting Text for Table S1

The following descriptions are based on the results of the Wilcoxon signed-rank tests, as our primary goal was to establish the relative ordering of stimulus groups in terms of perceived dynamism. Chi-square tests are reported in the table to provide additional information about differences in response distributions.

The first set of comparisons focused on stimuli with equal-sized elements combining four possible global orientations (0°, 90°, 45°, 135°) with two local edge orientations (cardinal 0–90°; oblique 45–135°). Several systematic effects emerged:

- First, when local edges were oblique, globally oblique configurations were rated as more dynamic than their globally cardinal counterparts [1.1]; the same pattern emerged when local edges were cardinal [1.2], indicating a strong and consistent influence of global orientation on perceived dynamism independently of local edge orientation.
- We then considered the effect of local edge orientation while keeping global orientation constant. Under a global cardinal layout, stimuli with oblique local edges were judged as more dynamic than those with cardinal edges [1.3]. Under a global oblique configuration, however, the opposite

pattern emerged: stimuli with cardinal local edges were rated as more dynamic than those with oblique edges [1.4]. However, this inversion likely reflects differences in the overall spatial dispersion of the configuration rather than a reversal of the contribution of local orientation, as oblique global layouts produce a more extended arrangement of cardinal-edged elements.

- We next examined contrasts in which both global and local orientations differed. Globally oblique configurations were more dynamic than globally cardinal ones even when the oblique configuration contained cardinal-edged (square) elements [1.5], demonstrating that the global layout alone can robustly elicit dynamism. Moreover, stimuli in which both global and local orientations were oblique (diamond-shaped elements arranged along an oblique axis) were substantially more dynamic than globally cardinal configurations composed of square elements [1.6].
- To isolate the effect of spatial dispersion, we compared stimuli with more widely dispersed versus more compact element distributions. Configurations with greater spatial dispersion were consistently judged as more dynamic [1.7], suggesting that dispersion acts as an additional visual cue enhancing perceived motion.
- We also compared horizontal and vertical global orientations. For stimuli with oblique local edges, horizontal configurations were perceived as less dynamic than vertical ones [1.8], whereas no significant difference emerged for stimuli with cardinal local edges [1.9], indicating an interaction between global orientation and local structure.
- Finally, comparing the two oblique global orientations (45° vs. 135°) revealed no significant differences in perceived dynamism either when local orientation is cardinal [1.10] or oblique [1.11], suggesting isotropy within the oblique axis.

The second set of comparisons examined stimuli with equal-sized elements combining the same four global orientations (0°, 90°, 45°, 135°) with two additional oblique local-edge pairs (30–150° and 105–165° for "fat" diamonds; 60–120° and 15–75° for "thin" diamonds). Results show that:

- In both cases, globally oblique configurations were again rated as more dynamic than globally cardinal ones [2.1, 2.2], confirming the dominant role of global orientation.
- As in the first set, horizontal configurations were judged as less dynamic than vertical ones [2.3, 2.4].
- Finally, to test whether the spacing effect observed in Set 1 generalized across different element shapes, we compared stimuli containing "fat" versus "thin" diamonds. The latter, which also exhibited greater spacing between neighboring elements, were perceived as more dynamic [2.5], indicating that increased separation between elements contributes to perceived dynamism across different shape configurations.

The third set focused on stimuli with equal-sized, cardinal-edged (square) elements arranged along two other oblique global orientations (30°/150° vs. 60°/120°):

- A significant difference emerged between these orientation pairs [3.1], indicating that even relatively small variations within the oblique range modulate perceived dynamism, with orientations closer to the horizontal axis being judged as less dynamic than those closer to the vertical axis.

Moving to stimuli with varying-size elements, the fourth set comprised horizontal (0°) and vertical (90°) configurations across four oblique local-edge orientations (0–90°, 30–150°, 45–135°, 60–120°). We found:

- No differences emerged between leftward and rightward configurations once global and local orientations were fixed [4.1].
- In contrast, upward configurations were rated as significantly more dynamic than downward configurations [4.2].

The fifth set tested stimuli arranged along all possible mirror-symmetric global oblique orientations (30°, 45°, 60°, and 150°, 135°, and 120°) combined with four oblique local-edge orientations (0–90°, 15–75°, 45–135°, and 105–165°). Here:

- No differences emerged between ascending and descending variants [5.1], indicating that alignment direction does not contribute to perceived dynamism once global and local orientations are fixed.

The last set comprised all equal-size stimuli with four possible global orientations (0°, 90°, 45°, 135°) and their corresponding varying-size counterparts. We found that:

- Across all global orientations, stimuli containing elements that varied systematically in size along the configuration axis – whether increasing or decreasing in the horizontal [6.1], vertical [6.2], or diagonal directions [6.3–6.4] – were rated as more dynamic than their equal-size counterparts. These results suggest that size gradients constitute an additional source of perceived dynamism.

**Table S2.** Stimulus pairs used to test the effect of global orientation.

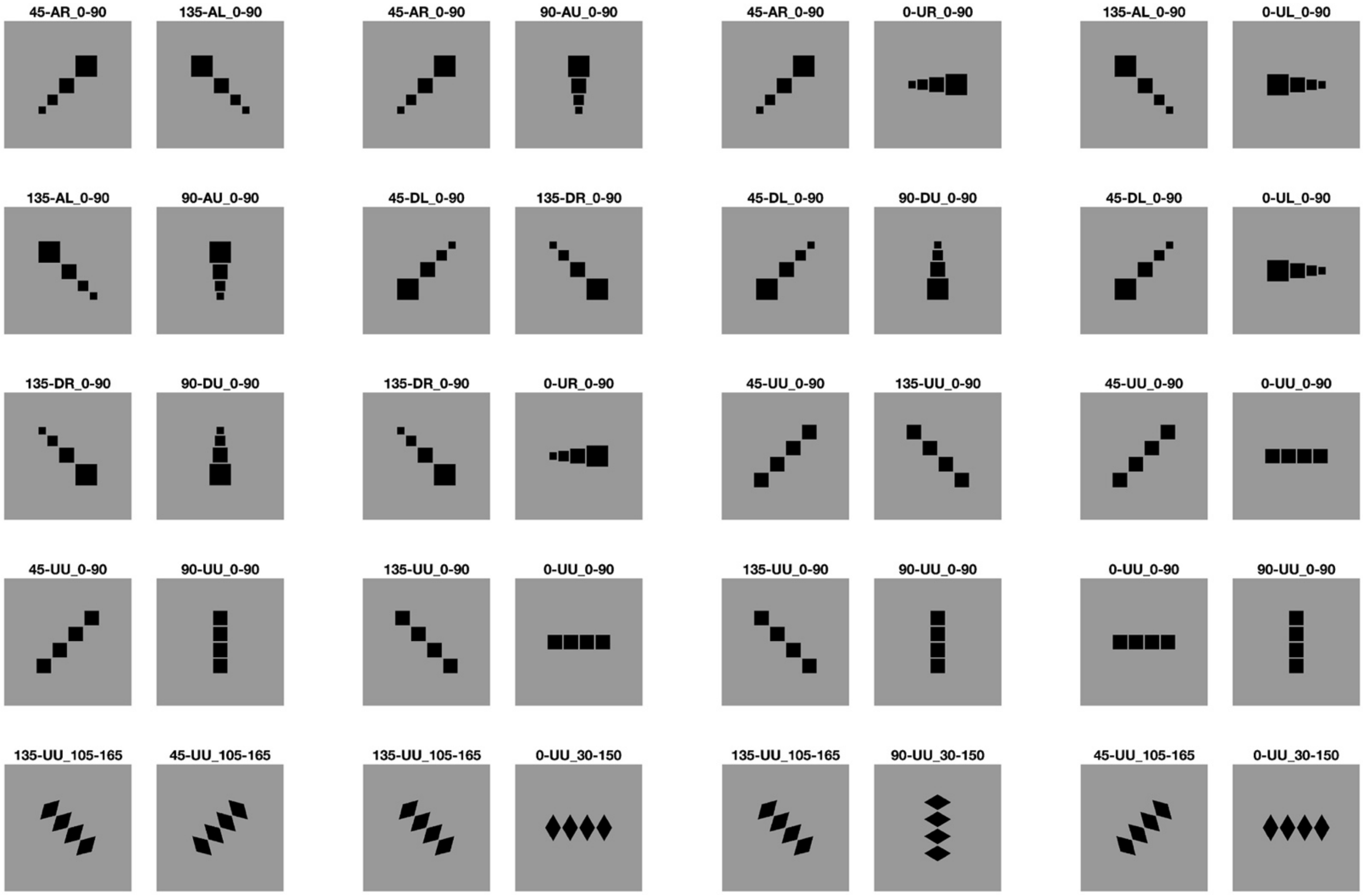

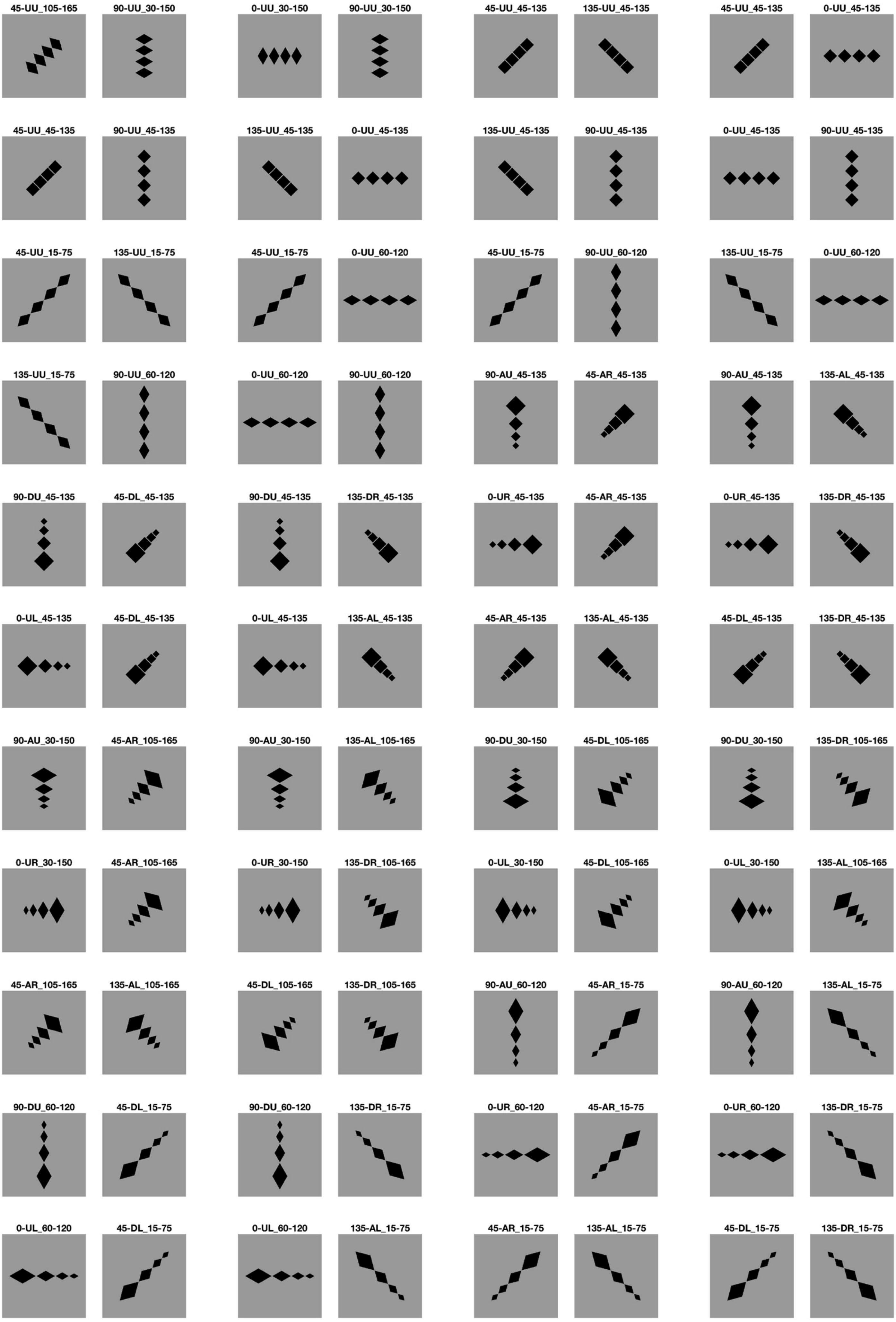
45-UU_105-165
90-UU_30-150
0-UU_30-150
90-UU_30-150
45-UU_45-135
135-UU_45-135
45-UU_45-135
0-UU_45-135
45-UU_45-135
90-UU_45-135
135-UU_45-135
0-UU_45-135
135-UU_45-135
90-UU_45-135
0-UU_45-135
90-UU_45-135
45-UU_15-75
135-UU_15-75
45-UU_15-75
0-UU_60-120
45-UU_15-75
90-UU_60-120
135-UU_15-75
0-UU_60-120
135-UU_15-75
90-UU_60-120
0-UU_60-120
90-UU_60-120
90-AU_45-135
45-AR_45-135
90-AU_45-135
135-AL_45-135
90-DU_45-135
45-DL_45-135
90-DU_45-135
135-DR_45-135
0-UR_45-135
45-AR_45-135
0-UR_45-135
135-DR_45-135
0-UL_45-135
45-DL_45-135
0-UL_45-135
135-AL_45-135
45-AR_45-135
135-AL_45-135
45-DL_45-135
135-DR_45-135
90-AU_30-150
45-AR_105-165
90-AU_30-150
135-AL_105-165
90-DU_30-150
45-DL_105-165
90-DU_30-150
135-DR_105-165
0-UR_30-150
45-AR_105-165
0-UR_30-150
135-DR_105-165
0-UL_30-150
45-DL_105-165
0-UL_30-150
135-AL_105-165
45-AR_105-165
135-AL_105-165
45-DL_105-165
135-DR_105-165
90-AU_60-120
45-AR_15-75
90-AU_60-120
135-AL_15-75
90-DU_60-120
45-DL_15-75
90-DU_60-120
135-DR_15-75
0-UR_60-120
45-AR_15-75
0-UR_60-120
135-DR_15-75
0-UL_60-120
45-DL_15-75
0-UL_60-120
135-AL_15-75
45-AR_15-75
135-AL_15-75
45-DL_15-75
135-DR_15-75

**Table S3.** Stimulus pairs used to test the effect of local orientation.

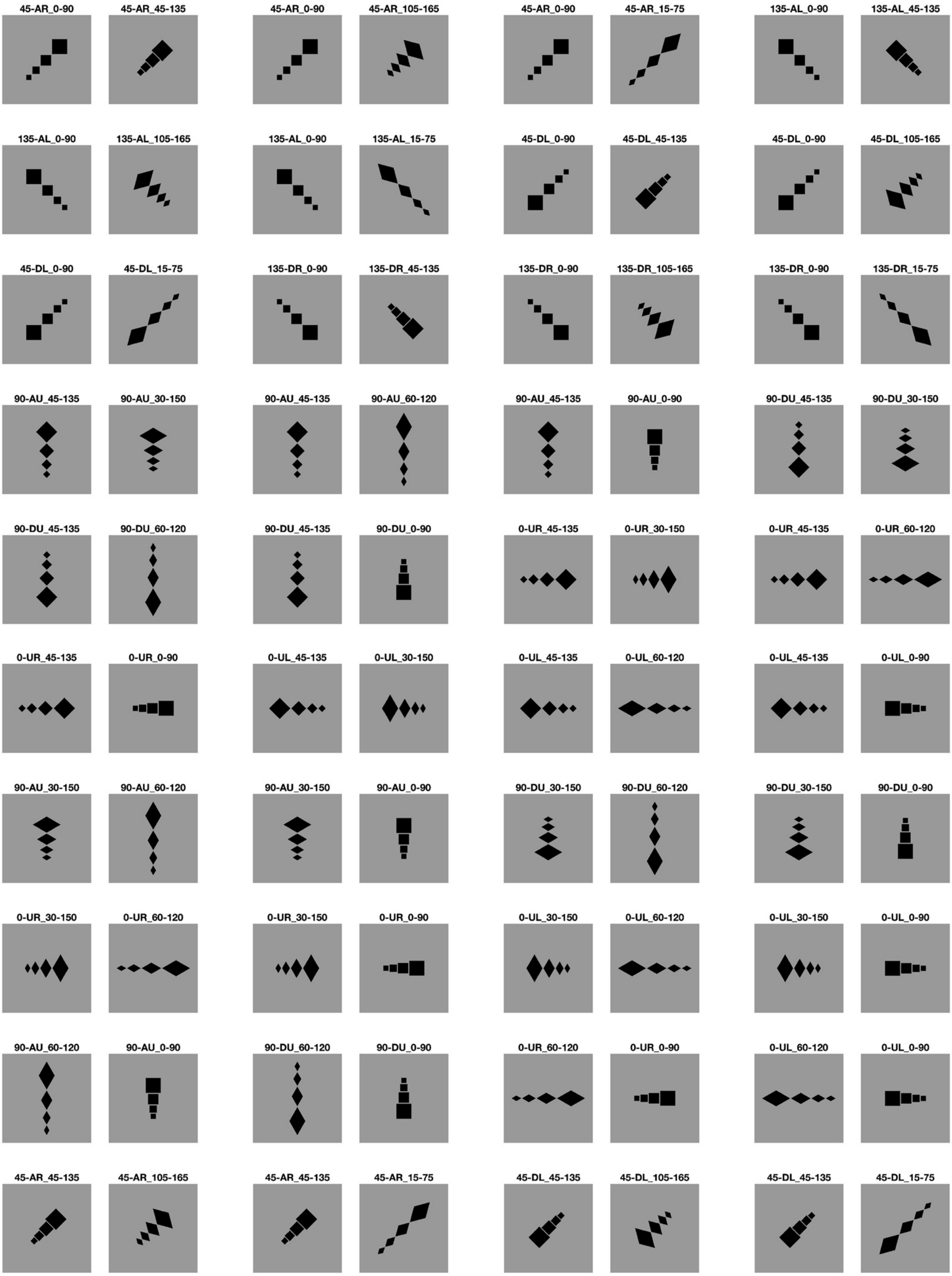

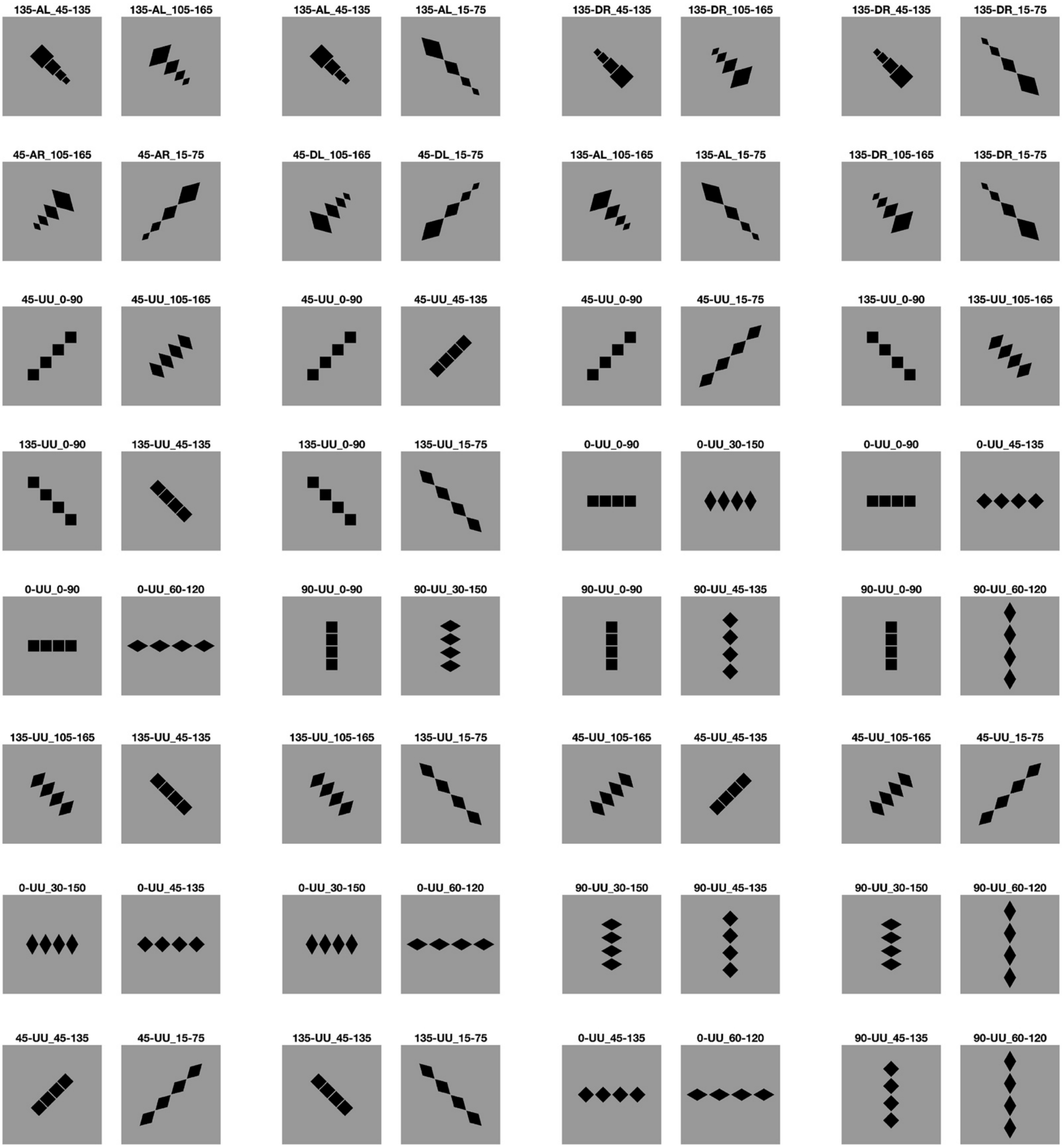

135-AL_45-135
135-AL_105-165
135-AL_45-135
135-AL_15-75
135-DR_45-135
135-DR_105-165
135-DR_45-135
135-DR_15-75
45-AR_105-165
45-AR_15-75
45-DL_105-165
45-DL_15-75
135-AL_105-165
135-AL_15-75
135-DR_105-165
135-DR_15-75
45-UU_0-90
45-UU_105-165
45-UU_0-90
45-UU_45-135
45-UU_0-90
45-UU_15-75
135-UU_0-90
135-UU_105-165
135-UU_0-90
135-UU_45-135
135-UU_0-90
135-UU_15-75
0-UU_0-90
0-UU_30-150
0-UU_0-90
0-UU_45-135
0-UU_0-90
0-UU_60-120
90-UU_0-90
90-UU_30-150
90-UU_0-90
90-UU_45-135
90-UU_0-90
90-UU_60-120
135-UU_105-165
135-UU_45-135
135-UU_105-165
135-UU_15-75
45-UU_105-165
45-UU_45-135
45-UU_105-165
45-UU_15-75
0-UU_30-150
0-UU_45-135
0-UU_30-150
0-UU_60-120
90-UU_30-150
90-UU_45-135
90-UU_30-150
90-UU_60-120
45-UU_45-135
45-UU_15-75
135-UU_45-135
135-UU_15-75
0-UU_45-135
0-UU_60-120
90-UU_45-135
90-UU_60-120

**Table S4.** Stimulus pairs used to test the effect of element size variation.

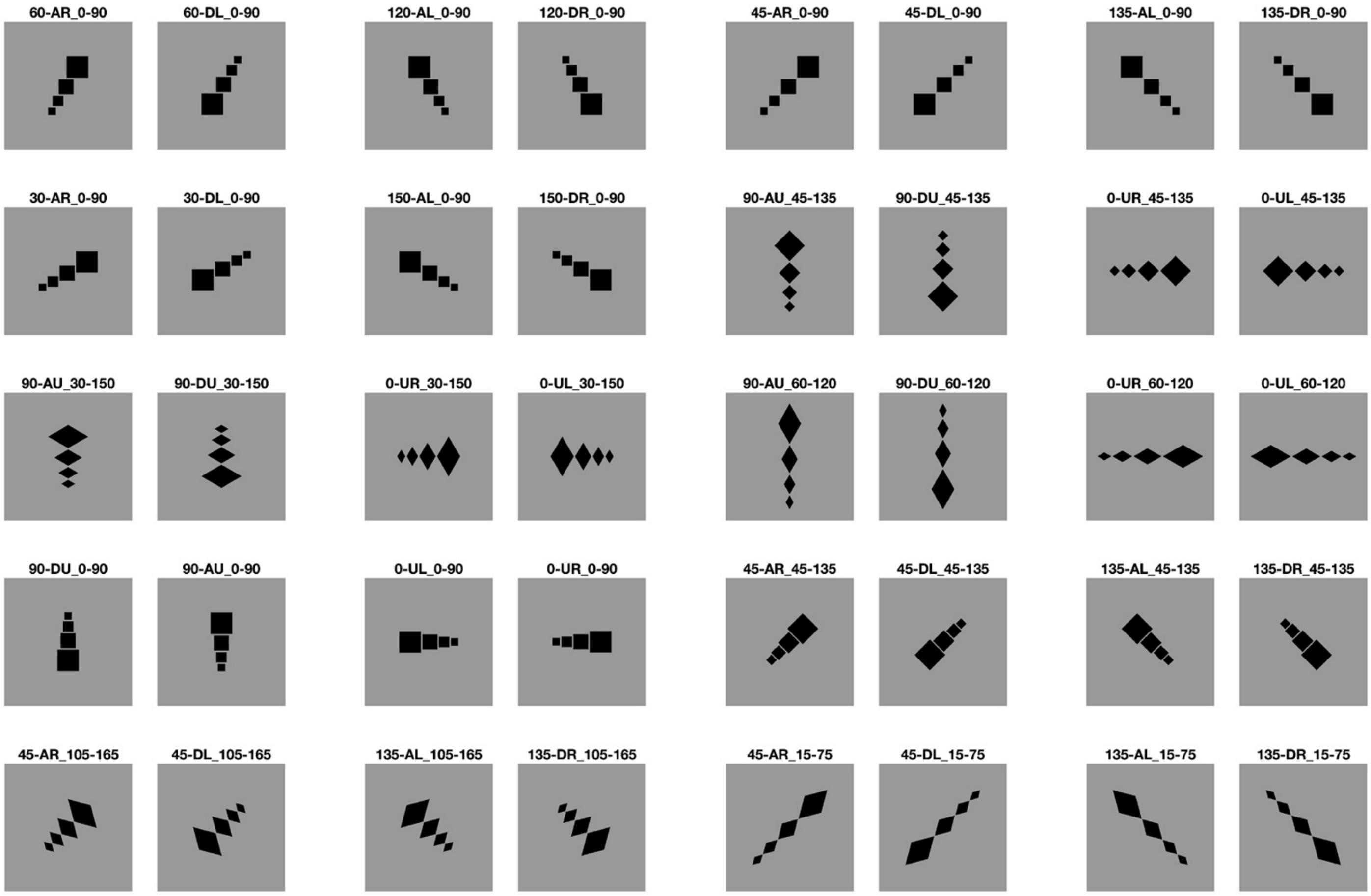

**Table S5.** Stimulus pairs used to test the effect of the interaction between global configuration orientation and local edge orientation.

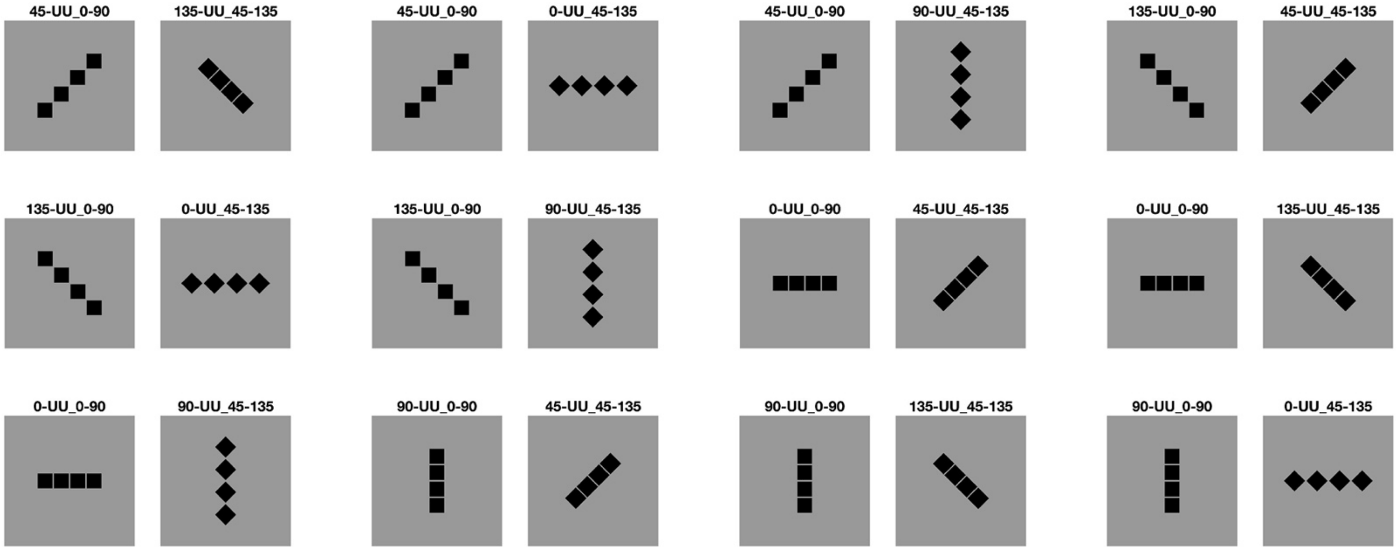

**Table S6. Experiment 1: Pairwise differences in mean dynamism ratings.** Matrix reporting the differences between the mean dynamism ratings of all possible pairs of stimuli.

| | 60-AR_0-90 | 120-AL_0-90 | 60-DL_0-90 | 120-DR_0-90 | 45-AR_0-90 | 135-AL_0-90 | 45-DL_0-90 | 135-DR_0-90 | 30-AR_0-90 | 150-AL_0-90 | 30-DL_0-90 | 150-DR_0-90 | 90-AU_45-135 | 90-DU_45-135 | 0-UR_45-135 | 0-UL_45-135 | 90-AU_30-150 | 90-DU_30-150 | 0-UR_30-150 | 0-UL_30-150 |
|---|---|---|---|---|---|---|---|---|---|---|---|---|---|---|---|---|---|---|---|---|
| **60-AR_0-90** | | 0.02 | -0.04 | -0.14 | -0.12 | -0.08 | -0.14 | -0.03 | -0.09 | -0.09 | -0.07 | 0.02 | -0.09 | 0.00 | 0.18 | 0.17 | -0.54 | -0.47 | -0.32 | -0.31 |
| **120-AL_0-90** | | | -0.06 | -0.16 | -0.14 | -0.10 | -0.16 | -0.05 | -0.11 | -0.11 | -0.09 | 0.00 | -0.11 | -0.02 | 0.16 | 0.15 | -0.56 | -0.49 | -0.34 | -0.33 |
| **60-DL_0-90** | | | | -0.10 | -0.08 | -0.04 | -0.10 | 0.01 | -0.05 | -0.05 | -0.03 | 0.06 | -0.05 | 0.04 | 0.22 | 0.21 | -0.50 | -0.43 | -0.28 | -0.27 |
| **120-DR_0-90** | | | | | 0.02 | 0.06 | 0.00 | 0.11 | 0.05 | 0.05 | 0.07 | 0.16 | 0.05 | 0.14 | 0.32 | 0.31 | -0.40 | -0.33 | -0.18 | -0.17 |
| **45-AR_0-90** | | | | | | 0.04 | -0.02 | 0.09 | 0.03 | 0.03 | 0.05 | 0.14 | 0.03 | 0.12 | 0.30 | 0.29 | -0.42 | -0.35 | -0.20 | -0.19 |
| **135-AL_0-90** | | | | | | | -0.06 | 0.05 | -0.01 | -0.01 | 0.01 | 0.10 | -0.01 | 0.08 | 0.26 | 0.25 | -0.46 | -0.39 | -0.24 | -0.23 |
| **45-DL_0-90** | | | | | | | | 0.11 | 0.05 | 0.05 | 0.07 | 0.16 | 0.05 | 0.14 | 0.32 | 0.31 | -0.40 | -0.33 | -0.18 | -0.17 |
| **135-DR_0-90** | | | | | | | | | -0.06 | -0.06 | -0.04 | 0.05 | -0.06 | 0.03 | 0.21 | 0.20 | -0.51 | -0.44 | -0.29 | -0.28 |
| **30-AR_0-90** | | | | | | | | | | 0.00 | 0.02 | 0.11 | 0.00 | 0.09 | 0.27 | 0.26 | -0.45 | -0.38 | -0.23 | -0.22 |
| **150-AL_0-90** | | | | | | | | | | | 0.02 | 0.11 | 0.00 | 0.09 | 0.27 | 0.26 | -0.45 | -0.38 | -0.23 | -0.22 |
| **30-DL_0-90** | | | | | | | | | | | | 0.09 | -0.02 | 0.07 | 0.25 | 0.24 | -0.47 | -0.40 | -0.25 | -0.24 |
| **150-DR_0-90** | | | | | | | | | | | | | -0.11 | -0.02 | 0.16 | 0.15 | -0.56 | -0.49 | -0.34 | -0.33 |
| **90-AU_45-135** | | | | | | | | | | | | | | 0.09 | 0.27 | 0.26 | -0.45 | -0.38 | -0.23 | -0.22 |
| **90-DU_45-135** | | | | | | | | | | | | | | | 0.18 | 0.17 | -0.54 | -0.47 | -0.32 | -0.31 |
| **0-UR_45-135** | | | | | | | | | | | | | | | | -0.01 | -0.72 | -0.65 | -0.50 | -0.49 |
| **0-UL_45-135** | | | | | | | | | | | | | | | | | -0.71 | -0.64 | -0.49 | -0.48 |
| **90-AU_30-150** | | | | | | | | | | | | | | | | | | 0.07 | 0.22 | 0.23 |
| **90-DU_30-150** | | | | | | | | | | | | | | | | | | | 0.15 | 0.16 |
| **0-UR_30-150** | | | | | | | | | | | | | | | | | | | | 0.01 |

| | 90-AU_60-120 | 90-DU_60-120 | 0-UR_60-120 | 0-UL_60-120 | 90-DU_0-90 | 0-UL_0-90 | 90-AU_0-90 | 0-UR_0-90 | 45-AR_45-135 | 45-DL_45-135 | 135-AL_45-135 | 135-DR_45-135 | 45-AR_105-165 | 45-DL_105-165 | 135-AL_105-165 | 135-DR_105-165 | 45-AR_15-75 | 45-DL_15-75 | 135-AL_15-75 | 135-DR_15-75 |
|---|---|---|---|---|---|---|---|---|---|---|---|---|---|---|---|---|---|---|---|---|
| **60-AR_0-90** | -0.22 | -0.21 | 0.05 | 0.10 | 0.39 | 0.22 | 0.14 | 0.18 | -0.19 | -0.06 | 0.04 | -0.02 | -0.64 | -0.64 | -0.72 | -0.68 | -0.44 | -0.44 | -0.48 | -0.57 |
| **120-AL_0-90** | -0.24 | -0.23 | 0.03 | 0.08 | 0.37 | 0.20 | 0.12 | 0.16 | -0.21 | -0.08 | 0.02 | -0.04 | -0.66 | -0.66 | -0.74 | -0.70 | -0.46 | -0.46 | -0.50 | -0.59 |
| **60-DL_0-90** | -0.18 | -0.17 | 0.09 | 0.14 | 0.43 | 0.26 | 0.18 | 0.22 | -0.15 | -0.02 | 0.08 | 0.02 | -0.60 | -0.60 | -0.68 | -0.64 | -0.40 | -0.40 | -0.44 | -0.53 |
| **120-DR_0-90** | -0.08 | -0.07 | 0.19 | 0.24 | 0.53 | 0.36 | 0.28 | 0.32 | -0.05 | 0.08 | 0.18 | 0.12 | -0.50 | -0.50 | -0.58 | -0.54 | -0.30 | -0.30 | -0.34 | -0.43 |
| **45-AR_0-90** | -0.10 | -0.09 | 0.17 | 0.22 | 0.51 | 0.34 | 0.26 | 0.30 | -0.07 | 0.06 | 0.16 | 0.10 | -0.52 | -0.52 | -0.60 | -0.56 | -0.32 | -0.32 | -0.36 | -0.45 |
| **135-AL_0-90** | -0.14 | -0.13 | 0.13 | 0.18 | 0.47 | 0.30 | 0.22 | 0.26 | -0.11 | 0.02 | 0.12 | 0.06 | -0.56 | -0.56 | -0.64 | -0.60 | -0.36 | -0.36 | -0.40 | -0.49 |
| **45-DL_0-90** | -0.08 | -0.07 | 0.19 | 0.24 | 0.53 | 0.36 | 0.28 | 0.32 | -0.05 | 0.08 | 0.18 | 0.12 | -0.50 | -0.50 | -0.58 | -0.54 | -0.30 | -0.30 | -0.34 | -0.43 |
| **135-DR_0-90** | -0.19 | -0.18 | 0.08 | 0.13 | 0.42 | 0.25 | 0.17 | 0.21 | -0.16 | -0.03 | 0.07 | 0.01 | -0.61 | -0.61 | -0.69 | -0.65 | -0.41 | -0.41 | -0.45 | -0.54 |
| **30-AR_0-90** | -0.13 | -0.12 | 0.14 | 0.19 | 0.48 | 0.31 | 0.23 | 0.27 | -0.10 | 0.03 | 0.13 | 0.07 | -0.55 | -0.55 | -0.63 | -0.59 | -0.35 | -0.35 | -0.39 | -0.48 |
| **150-AL_0-90** | -0.13 | -0.12 | 0.14 | 0.19 | 0.48 | 0.31 | 0.23 | 0.27 | -0.10 | 0.03 | 0.13 | 0.07 | -0.55 | -0.55 | -0.63 | -0.59 | -0.35 | -0.35 | -0.39 | -0.48 |
| **30-DL_0-90** | -0.15 | -0.14 | 0.12 | 0.17 | 0.46 | 0.29 | 0.21 | 0.25 | -0.12 | 0.01 | 0.11 | 0.05 | -0.57 | -0.57 | -0.65 | -0.61 | -0.37 | -0.37 | -0.41 | -0.50 |
| **150-DR_0-90** | -0.24 | -0.23 | 0.03 | 0.08 | 0.37 | 0.20 | 0.12 | 0.16 | -0.21 | -0.08 | 0.02 | -0.04 | -0.66 | -0.66 | -0.74 | -0.70 | -0.46 | -0.46 | -0.50 | -0.59 |
| **90-AU_45-135** | -0.13 | -0.12 | 0.14 | 0.19 | 0.48 | 0.31 | 0.23 | 0.27 | -0.10 | 0.03 | 0.13 | 0.07 | -0.55 | -0.55 | -0.63 | -0.59 | -0.35 | -0.35 | -0.39 | -0.48 |
| **90-DU_45-135** | -0.22 | -0.21 | 0.05 | 0.10 | 0.39 | 0.22 | 0.14 | 0.18 | -0.19 | -0.06 | 0.04 | -0.02 | -0.64 | -0.64 | -0.72 | -0.68 | -0.44 | -0.44 | -0.48 | -0.57 |
| **0-UR_45-135** | -0.40 | -0.39 | -0.13 | -0.08 | 0.21 | 0.04 | -0.04 | 0.00 | -0.37 | -0.24 | -0.14 | -0.20 | -0.82 | -0.82 | -0.90 | -0.86 | -0.62 | -0.62 | -0.66 | -0.75 |
| **0-UL_45-135** | -0.39 | -0.38 | -0.12 | -0.07 | 0.22 | 0.05 | -0.03 | 0.01 | -0.36 | -0.23 | -0.13 | -0.19 | -0.81 | -0.81 | -0.89 | -0.85 | -0.61 | -0.61 | -0.65 | -0.74 |
| **90-AU_30-150** | 0.32 | 0.33 | 0.59 | 0.64 | 0.93 | 0.76 | 0.68 | 0.72 | 0.35 | 0.48 | 0.58 | 0.52 | -0.10 | -0.10 | -0.18 | -0.14 | 0.10 | 0.10 | 0.06 | -0.03 |
| **90-DU_30-150** | 0.25 | 0.26 | 0.52 | 0.57 | 0.86 | 0.69 | 0.61 | 0.65 | 0.28 | 0.41 | 0.51 | 0.45 | -0.17 | -0.17 | -0.25 | -0.21 | 0.03 | 0.03 | -0.01 | -0.10 |
| **0-UR_30-150** | 0.10 | 0.11 | 0.37 | 0.42 | 0.71 | 0.54 | 0.46 | 0.50 | 0.13 | 0.26 | 0.36 | 0.30 | -0.32 | -0.32 | -0.40 | -0.36 | -0.12 | -0.12 | -0.16 | -0.25 |
| **0-UL_30-150** | 0.09 | 0.10 | 0.36 | 0.41 | 0.70 | 0.53 | 0.45 | 0.49 | 0.12 | 0.25 | 0.35 | 0.29 | -0.33 | -0.33 | -0.41 | -0.37 | -0.13 | -0.13 | -0.17 | -0.26 |
| **90-AU_60-120** | | 0.01 | 0.27 | 0.32 | 0.61 | 0.44 | 0.36 | 0.40 | 0.03 | 0.16 | 0.26 | 0.20 | -0.42 | -0.42 | -0.50 | -0.46 | -0.22 | -0.22 | -0.26 | -0.35 |
| **90-DU_60-120** | | | 0.26 | 0.31 | 0.60 | 0.43 | 0.35 | 0.39 | 0.02 | 0.15 | 0.25 | 0.19 | -0.43 | -0.43 | -0.51 | -0.47 | -0.23 | -0.23 | -0.27 | -0.36 |
| **0-UR_60-120** | | | | 0.05 | 0.34 | 0.17 | 0.09 | 0.13 | -0.24 | -0.11 | -0.01 | -0.07 | -0.69 | -0.69 | -0.77 | -0.73 | -0.49 | -0.49 | -0.53 | -0.62 |
| **0-UL_60-120** | | | | | 0.29 | 0.12 | 0.04 | 0.08 | -0.29 | -0.16 | -0.06 | -0.12 | -0.74 | -0.74 | -0.82 | -0.78 | -0.54 | -0.54 | -0.58 | -0.67 |
| **90-DU_0-90** | | | | | | -0.17 | -0.25 | -0.21 | -0.58 | -0.45 | -0.35 | -0.41 | -1.03 | -1.03 | -1.11 | -1.07 | -0.83 | -0.83 | -0.87 | -0.96 |
| **0-UL_0-90** | | | | | | | -0.08 | -0.04 | -0.41 | -0.28 | -0.18 | -0.24 | -0.86 | -0.86 | -0.94 | -0.90 | -0.66 | -0.66 | -0.70 | -0.79 |
| **90-AU_0-90** | | | | | | | | 0.04 | -0.33 | -0.20 | -0.10 | -0.16 | -0.78 | -0.78 | -0.86 | -0.82 | -0.58 | -0.58 | -0.62 | -0.71 |
| **0-UR_0-90** | | | | | | | | | -0.37 | -0.24 | -0.14 | -0.20 | -0.82 | -0.82 | -0.90 | -0.86 | -0.62 | -0.62 | -0.66 | -0.75 |

| | 30-UU_0-90 | 45-UU_0-90 | 120-UU_0-90 | 135-UU_0-90 | 150-UU_0-90 | 60-UU_0-90 | 0-UU_0-90 | 90-UU_0-90 | 135-UU_105-165 | 45-UU_105-165 | 0-UU_30-150 | 90-UU_30-150 | 45-UU_45-135 | 135-UU_45-135 | 0-UU_45-135 | 90-UU_45-135 | 45-UU_15-75 | 135-UU_15-75 | 0-UU_60-120 | 90-UU_60-120 |
|---|---|---|---|---|---|---|---|---|---|---|---|---|---|---|---|---|---|---|---|---|
| **60-AR_0-90** | 0.22 | 0.31 | 0.17 | 0.31 | 0.26 | 0.06 | 1.46 | 1.34 | -0.34 | -0.19 | 0.74 | 0.22 | 0.71 | 0.55 | 1.03 | 0.68 | -0.13 | -0.11 | 0.70 | 0.43 |
| **120-AL_0-90** | 0.20 | 0.29 | 0.15 | 0.29 | 0.24 | 0.04 | 1.44 | 1.32 | -0.36 | -0.21 | 0.72 | 0.20 | 0.69 | 0.53 | 1.01 | 0.66 | -0.15 | -0.13 | 0.68 | 0.41 |
| **60-DL_0-90** | 0.26 | 0.35 | 0.21 | 0.35 | 0.30 | 0.10 | 1.50 | 1.38 | -0.30 | -0.15 | 0.78 | 0.26 | 0.75 | 0.59 | 1.07 | 0.72 | -0.09 | -0.07 | 0.74 | 0.47 |
| **120-DR_0-90** | 0.36 | 0.45 | 0.31 | 0.45 | 0.40 | 0.20 | 1.60 | 1.48 | -0.20 | -0.05 | 0.88 | 0.36 | 0.85 | 0.69 | 1.17 | 0.82 | 0.01 | 0.03 | 0.84 | 0.57 |
| **45-AR_0-90** | 0.34 | 0.43 | 0.29 | 0.43 | 0.38 | 0.18 | 1.58 | 1.46 | -0.22 | -0.07 | 0.86 | 0.34 | 0.83 | 0.67 | 1.15 | 0.80 | -0.01 | 0.01 | 0.82 | 0.55 |
| **135-AL_0-90** | 0.30 | 0.39 | 0.25 | 0.39 | 0.34 | 0.14 | 1.54 | 1.42 | -0.26 | -0.11 | 0.82 | 0.30 | 0.79 | 0.63 | 1.11 | 0.76 | -0.05 | -0.03 | 0.78 | 0.51 |
| **45-DL_0-90** | 0.36 | 0.45 | 0.31 | 0.45 | 0.40 | 0.20 | 1.60 | 1.48 | -0.20 | -0.05 | 0.88 | 0.36 | 0.85 | 0.69 | 1.17 | 0.82 | 0.01 | 0.03 | 0.84 | 0.57 |
| **135-DR_0-90** | 0.25 | 0.34 | 0.20 | 0.34 | 0.29 | 0.09 | 1.49 | 1.37 | -0.31 | -0.16 | 0.77 | 0.25 | 0.74 | 0.58 | 1.06 | 0.71 | -0.10 | -0.08 | 0.73 | 0.46 |
| **30-AR_0-90** | 0.31 | 0.40 | 0.26 | 0.40 | 0.35 | 0.15 | 1.55 | 1.43 | -0.25 | -0.10 | 0.83 | 0.31 | 0.80 | 0.64 | 1.12 | 0.77 | -0.04 | -0.02 | 0.79 | 0.52 |
| **150-AL_0-90** | 0.31 | 0.40 | 0.26 | 0.40 | 0.35 | 0.15 | 1.55 | 1.43 | -0.25 | -0.10 | 0.83 | 0.31 | 0.80 | 0.64 | 1.12 | 0.77 | -0.04 | -0.02 | 0.79 | 0.52 |
| **30-DL_0-90** | 0.29 | 0.38 | 0.24 | 0.38 | 0.33 | 0.13 | 1.53 | 1.41 | -0.27 | -0.12 | 0.81 | 0.29 | 0.78 | 0.62 | 1.10 | 0.75 | -0.06 | -0.04 | 0.77 | 0.50 |
| **150-DR_0-90** | 0.20 | 0.29 | 0.15 | 0.29 | 0.24 | 0.04 | 1.44 | 1.32 | -0.36 | -0.21 | 0.72 | 0.20 | 0.69 | 0.53 | 1.01 | 0.66 | -0.15 | -0.13 | 0.68 | 0.41 |
| **90-AU_45-135** | 0.31 | 0.40 | 0.26 | 0.40 | 0.35 | 0.15 | 1.55 | 1.43 | -0.25 | -0.10 | 0.83 | 0.31 | 0.80 | 0.64 | 1.12 | 0.77 | -0.04 | -0.02 | 0.79 | 0.52 |
| **90-DU_45-135** | 0.22 | 0.31 | 0.17 | 0.31 | 0.26 | 0.06 | 1.46 | 1.34 | -0.34 | -0.19 | 0.74 | 0.22 | 0.71 | 0.55 | 1.03 | 0.68 | -0.13 | -0.11 | 0.70 | 0.43 |
| **0-UR_45-135** | 0.04 | 0.13 | -0.01 | 0.13 | 0.08 | -0.12 | 1.28 | 1.16 | -0.52 | -0.37 | 0.56 | 0.04 | 0.53 | 0.37 | 0.85 | 0.50 | -0.31 | -0.29 | 0.52 | 0.25 |
| **0-UL_45-135** | 0.05 | 0.14 | 0.00 | 0.14 | 0.09 | -0.11 | 1.29 | 1.17 | -0.51 | -0.36 | 0.57 | 0.05 | 0.54 | 0.38 | 0.86 | 0.51 | -0.30 | -0.28 | 0.53 | 0.26 |
| **90-AU_30-150** | 0.76 | 0.85 | 0.71 | 0.85 | 0.80 | 0.60 | 2.00 | 1.88 | 0.20 | 0.35 | 1.28 | 0.76 | 1.25 | 1.09 | 1.57 | 1.22 | 0.41 | 0.43 | 1.24 | 0.97 |
| **90-DU_30-150** | 0.69 | 0.78 | 0.64 | 0.78 | 0.73 | 0.53 | 1.93 | 1.81 | 0.13 | 0.28 | 1.21 | 0.69 | 1.18 | 1.02 | 1.50 | 1.15 | 0.34 | 0.36 | 1.17 | 0.90 |
| **0-UR_30-150** | 0.54 | 0.63 | 0.49 | 0.63 | 0.58 | 0.38 | 1.78 | 1.66 | -0.02 | 0.13 | 1.06 | 0.54 | 1.03 | 0.87 | 1.35 | 1.00 | 0.19 | 0.21 | 1.02 | 0.75 |
| **0-UL_30-150** | 0.53 | 0.62 | 0.48 | 0.62 | 0.57 | 0.37 | 1.77 | 1.65 | -0.03 | 0.12 | 1.05 | 0.53 | 1.02 | 0.86 | 1.34 | 0.99 | 0.18 | 0.20 | 1.01 | 0.74 |
| **90-AU_60-120** | 0.44 | 0.53 | 0.39 | 0.53 | 0.48 | 0.28 | 1.68 | 1.56 | -0.12 | 0.03 | 0.96 | 0.44 | 0.93 | 0.77 | 1.25 | 0.90 | 0.09 | 0.11 | 0.92 | 0.65 |
| **90-DU_60-120** | 0.43 | 0.52 | 0.38 | 0.52 | 0.47 | 0.27 | 1.67 | 1.55 | -0.13 | 0.02 | 0.95 | 0.43 | 0.92 | 0.76 | 1.24 | 0.89 | 0.08 | 0.10 | 0.91 | 0.64 |
| **0-UR_60-120** | 0.17 | 0.26 | 0.12 | 0.26 | 0.21 | 0.01 | 1.41 | 1.29 | -0.39 | -0.24 | 0.69 | 0.17 | 0.66 | 0.50 | 0.98 | 0.63 | -0.18 | -0.16 | 0.65 | 0.38 |
| **0-UL_60-120** | 0.12 | 0.21 | 0.07 | 0.21 | 0.16 | -0.04 | 1.36 | 1.24 | -0.44 | -0.29 | 0.64 | 0.12 | 0.61 | 0.45 | 0.93 | 0.58 | -0.23 | -0.21 | 0.60 | 0.33 |
| **90-DU_0-90** | -0.17 | -0.08 | -0.22 | -0.08 | -0.13 | -0.33 | 1.07 | 0.95 | -0.73 | -0.58 | 0.35 | -0.17 | 0.32 | 0.16 | 0.64 | 0.29 | -0.52 | -0.50 | 0.31 | 0.04 |
| **0-UL_0-90** | 0.00 | 0.09 | -0.05 | 0.09 | 0.04 | -0.16 | 1.24 | 1.12 | -0.56 | -0.41 | 0.52 | 0.00 | 0.49 | 0.33 | 0.81 | 0.46 | -0.35 | -0.33 | 0.48 | 0.21 |
| **90-AU_0-90** | 0.08 | 0.17 | 0.03 | 0.17 | 0.12 | -0.08 | 1.32 | 1.20 | -0.48 | -0.33 | 0.60 | 0.08 | 0.57 | 0.41 | 0.89 | 0.54 | -0.27 | -0.25 | 0.56 | 0.29 |
| **0-UR_0-90** | 0.04 | 0.13 | -0.01 | 0.13 | 0.08 | -0.12 | 1.28 | 1.16 | -0.52 | -0.37 | 0.56 | 0.04 | 0.53 | 0.37 | 0.85 | 0.50 | -0.31 | -0.29 | 0.52 | 0.25 |

| | | | | | | | | | | | | | | | | | | | | | | | | | | | | | | | |
|---|---|---|---|---|---|---|---|---|---|---|---|---|---|---|---|---|---|---|---|---|---|---|---|---|---|---|---|---|---|---|---|
| 0.62 | 0.49 | 0.39 | 0.45 | 1.07 | 1.07 | 1.15 | 1.11 | 0.87 | 0.87 | 0.91 | 1.00 | 0.21 | 0.12 | 0.26 | 0.12 | 0.17 | 0.37 | -1.03 | -0.91 | 0.77 | 0.62 | -0.31 | 0.21 | -0.28 | -0.12 | -0.60 | -0.25 | 0.56 | 0.54 | -0.27 | |
| 0.89 | 0.76 | 0.66 | 0.72 | 1.34 | 1.34 | 1.42 | 1.38 | 1.14 | 1.14 | 1.18 | 1.27 | 0.48 | 0.39 | 0.53 | 0.39 | 0.44 | 0.64 | -0.76 | -0.64 | 1.04 | 0.89 | -0.04 | 0.48 | -0.01 | 0.15 | -0.33 | 0.02 | 0.83 | 0.81 | | |
| 0.08 | -0.05 | -0.15 | -0.09 | 0.53 | 0.53 | 0.61 | 0.57 | 0.33 | 0.33 | 0.37 | 0.46 | -0.33 | -0.42 | -0.28 | -0.42 | -0.37 | -0.17 | -1.57 | -1.45 | 0.23 | 0.08 | -0.85 | -0.33 | -0.82 | -0.66 | -1.14 | -0.79 | 0.02 | | | |
| 0.06 | -0.07 | -0.17 | -0.11 | 0.51 | 0.51 | 0.59 | 0.55 | 0.31 | 0.31 | 0.35 | 0.44 | -0.35 | -0.44 | -0.30 | -0.44 | -0.39 | -0.19 | -1.59 | -1.47 | 0.21 | 0.06 | -0.87 | -0.35 | -0.84 | -0.68 | -1.16 | -0.81 | | | | |
| 0.87 | 0.74 | 0.64 | 0.70 | 1.32 | 1.32 | 1.40 | 1.36 | 1.12 | 1.12 | 1.16 | 1.25 | 0.46 | 0.37 | 0.51 | 0.37 | 0.42 | 0.62 | -0.78 | -0.66 | 1.02 | 0.87 | -0.06 | 0.46 | -0.03 | 0.13 | -0.35 | | | | | |
| 1.22 | 1.09 | 0.99 | 1.05 | 1.67 | 1.67 | 1.75 | 1.71 | 1.47 | 1.47 | 1.51 | 1.60 | 0.81 | 0.72 | 0.86 | 0.72 | 0.77 | 0.97 | -0.43 | -0.31 | 1.37 | 1.22 | 0.29 | 0.81 | 0.32 | 0.48 | | | | | | |
| 0.74 | 0.61 | 0.51 | 0.57 | 1.19 | 1.19 | 1.27 | 1.23 | 0.99 | 0.99 | 1.03 | 1.12 | 0.33 | 0.24 | 0.38 | 0.24 | 0.29 | 0.49 | -0.91 | -0.79 | 0.89 | 0.74 | -0.19 | 0.33 | -0.16 | | | | | | | |
| 0.90 | 0.77 | 0.67 | 0.73 | 1.35 | 1.35 | 1.43 | 1.39 | 1.15 | 1.15 | 1.19 | 1.28 | 0.49 | 0.40 | 0.54 | 0.40 | 0.45 | 0.65 | -0.75 | -0.63 | 1.05 | 0.90 | -0.03 | 0.49 | | | | | | | | |
| 0.41 | 0.28 | 0.18 | 0.24 | 0.86 | 0.86 | 0.94 | 0.90 | 0.66 | 0.66 | 0.70 | 0.79 | 0.00 | -0.09 | 0.05 | -0.09 | -0.04 | 0.16 | -1.24 | -1.12 | 0.56 | 0.41 | -0.52 | | | | | | | | | |
| 0.93 | 0.80 | 0.70 | 0.76 | 1.38 | 1.38 | 1.46 | 1.42 | 1.18 | 1.18 | 1.22 | 1.31 | 0.52 | 0.43 | 0.57 | 0.43 | 0.48 | 0.68 | -0.72 | -0.60 | 1.08 | 0.93 | | | | | | | | | | |
| 0.00 | -0.13 | -0.23 | -0.17 | 0.45 | 0.45 | 0.53 | 0.49 | 0.25 | 0.25 | 0.29 | 0.38 | -0.41 | -0.50 | -0.36 | -0.50 | -0.45 | -0.25 | -1.65 | -1.53 | 0.15 | | | | | | | | | | | |
| -0.15 | -0.28 | -0.38 | -0.32 | 0.30 | 0.30 | 0.38 | 0.34 | 0.10 | 0.10 | 0.14 | 0.23 | -0.56 | -0.65 | -0.51 | -0.65 | -0.60 | -0.40 | -1.80 | -1.68 | | | | | | | | | | | | |
| 1.53 | 1.40 | 1.30 | 1.36 | 1.98 | 1.98 | 2.06 | 2.02 | 1.78 | 1.78 | 1.82 | 1.91 | 1.12 | 1.03 | 1.17 | 1.03 | 1.08 | 1.28 | -0.12 | | | | | | | | | | | | | |
| 1.65 | 1.52 | 1.42 | 1.48 | 2.10 | 2.10 | 2.18 | 2.14 | 1.90 | 1.90 | 1.94 | 2.03 | 1.24 | 1.15 | 1.29 | 1.15 | 1.20 | 1.40 | | | | | | | | | | | | | | |
| 0.25 | 0.12 | 0.02 | 0.08 | 0.70 | 0.70 | 0.78 | 0.74 | 0.50 | 0.50 | 0.54 | 0.63 | -0.16 | -0.25 | -0.11 | -0.25 | -0.20 | | | | | | | | | | | | | | | |
| 0.45 | 0.32 | 0.22 | 0.28 | 0.90 | 0.90 | 0.98 | 0.94 | 0.70 | 0.70 | 0.74 | 0.83 | 0.04 | -0.05 | 0.09 | -0.05 | | | | | | | | | | | | | | | | |
| 0.50 | 0.37 | 0.27 | 0.33 | 0.95 | 0.95 | 1.03 | 0.99 | 0.75 | 0.75 | 0.79 | 0.88 | 0.09 | 0.00 | 0.14 | | | | | | | | | | | | | | | | | |
| 0.36 | 0.23 | 0.13 | 0.19 | 0.81 | 0.81 | 0.89 | 0.85 | 0.61 | 0.61 | 0.65 | 0.74 | -0.05 | -0.14 | | | | | | | | | | | | | | | | | | |
| 0.50 | 0.37 | 0.27 | 0.33 | 0.95 | 0.95 | 1.03 | 0.99 | 0.75 | 0.75 | 0.79 | 0.88 | 0.09 | | | | | | | | | | | | | | | | | | | |
| 0.41 | 0.28 | 0.18 | 0.24 | 0.86 | 0.86 | 0.94 | 0.90 | 0.66 | 0.66 | 0.70 | 0.79 | | | | | | | | | | | | | | | | | | | | |
| -0.38 | -0.51 | -0.61 | -0.55 | 0.07 | 0.07 | 0.15 | 0.11 | -0.13 | -0.13 | -0.09 | | | | | | | | | | | | | | | | | | | | | |
| -0.29 | -0.42 | -0.52 | -0.46 | 0.16 | 0.16 | 0.24 | 0.20 | -0.04 | -0.04 | | | | | | | | | | | | | | | | | | | | | | |
| -0.25 | -0.38 | -0.48 | -0.42 | 0.20 | 0.20 | 0.28 | 0.24 | 0.00 | | | | | | | | | | | | | | | | | | | | | | | |
| -0.25 | -0.38 | -0.48 | -0.42 | 0.20 | 0.20 | 0.28 | 0.24 | | | | | | | | | | | | | | | | | | | | | | | | |
| -0.49 | -0.62 | -0.72 | -0.66 | -0.04 | -0.04 | 0.04 | | | | | | | | | | | | | | | | | | | | | | | | | |
| -0.53 | -0.66 | -0.76 | -0.70 | -0.08 | -0.08 | | | | | | | | | | | | | | | | | | | | | | | | | | |
| -0.45 | -0.58 | -0.68 | -0.62 | 0.00 | | | | | | | | | | | | | | | | | | | | | | | | | | | |
| -0.45 | -0.58 | -0.68 | -0.62 | | | | | | | | | | | | | | | | | | | | | | | | | | | | |
| 0.17 | 0.04 | -0.06 | | | | | | | | | | | | | | | | | | | | | | | | | | | | | |
| 0.23 | 0.10 | | | | | | | | | | | | | | | | | | | | | | | | | | | | | | |
| 0.13 | | | | | | | | | | | | | | | | | | | | | | | | | | | | | | | |
| **45-AR_45-135** | **45-DL_45-135** | **135-AL_45-135** | **135-DR_45-135** | **45-AR_105-165** | **45-DL_105-165** | **135-AL_105-165** | **135-DR_105-165** | **45-AR_15-75** | **45-DL_15-75** | **135-AL_15-75** | **135-DR_15-75** | **30-UU_0-90** | **45-UU_0-90** | **120-UU_0-90** | **135-UU_0-90** | **150-UU_0-90** | **60-UU_0-90** | **0-UU_0-90** | **90-UU_0-90** | **135-UU_105-165** | **45-UU_105-165** | **0-UU_30-150** | **90-UU_30-150** | **45-UU_45-135** | **135-UU_45-135** | **0-UU_45-135** | **90-UU_45-135** | **45-UU_15-75** | **135-UU_15-75** | **0-UU_60-120** | **90-UU_60-120** |

**Table S7. Experiment 2: Mean pairwise dynamism scores.** Matrix reporting the mean pairwise dynamism scores for all stimulus pairs tested in Experiment 2.

| | 60-AR_0-90 | 120-AL_0-90 | 60-DL_0-90 | 120-DR_0-90 | 45-AR_0-90 | 135-AL_0-90 | 45-DL_0-90 | 135-DR_0-90 | 30-AR_0-90 | 150-AL_0-90 | 30-DL_0-90 | 150-DR_0-90 | 90-AU_45-135 | 90-DU_45-135 | 0-UR_45-135 | 0-UL_45-135 | 90-AU_30-150 | 90-DU_30-150 | 0-UR_30-150 | 0-UL_30-150 | 90-AU_60-120 | 90-DU_60-120 | 0-UR_60-120 | 0-UL_60-120 | 90-DU_0-90 | 0-UL_0-90 | 90-AU_0-90 | 0-UR_0-90 | 45-AR_45-135 | 45-DL_45-135 | 135-AL_45-135 | 135-DR_45-135 | 45-AR_105-165 | 45-DL_105-165 | 135-AL_105-165 | 135-DR_105-165 | 45-AR_15-75 | 45-DL_15-75 | 135-AL_15-75 | 135-DR_15-75 | 30-UU_0-90 | 45-UU_0-90 | 120-UU_0-90 | 135-UU_0-90 | 150-UU_0-90 | 60-UU_0-90 | 0-UU_0-90 | 90-UU_0-90 | 135-UU_105-165 | 45-UU_105-165 | 0-UU_30-150 | 90-UU_30-150 | 45-UU_45-135 | 135-UU_45-135 | 0-UU_45-135 | 90-UU_45-135 | 45-UU_15-75 | 135-UU_15-75 | 0-UU_60-120 | 90-UU_60-120 |
|---|---|---|---|---|---|---|---|---|---|---|---|---|---|---|---|---|---|---|---|---|---|---|---|---|---|---|---|---|---|---|---|---|---|---|---|---|---|---|---|---|---|---|---|---|---|---|---|---|---|---|---|---|---|---|---|---|---|---|---|---|
| **60-AR_0-90** | | | -0.07 | | | | | | | | | | | | | | | | | | | | | | | | | | | | | | | | | | | | | | | | | | | | | | | | | | | | | | | | | |
| **120-AL_0-90** | | | | 0.04 | | | | | | | | | | | | | | | | | | | | | | | | | | | | | | | | | | | | | | | | | | | | | | | | | | | | | | | | |
| **60-DL_0-90** | | | | | | | | | | | | | | | | | | | | | | | | | | | | | | | | | | | | | | | | | | | | | | | | | | | | | | | | | | | | |
| **120-DR_0-90** | | | | | | | | | | | | | | | | | | | | | | | | | | | | | | | | | | | | | | | | | | | | | | | | | | | | | | | | | | | | |
| **45-AR_0-90** | | | | | | -0.18 | -0.31 | | | | | | | | | | | | | | | | | | | | -0.78 | -0.78 | -0.68 | | | | 0.63 | | | | 0.95 | | | | | | | | | | | | | | | | | | | | | | | |
| **135-AL_0-90** | | | | | | | | 0.18 | | | | | | | | | | | | | | | | | | -0.83 | -0.82 | | | | -0.77 | | | | 0.71 | | | | 0.93 | | | | | | | | | | | | | | | | | | | | | |
| **45-DL_0-90** | | | | | | | | 0.06 | | | | | | | | | | | | | | | | | -0.88 | -0.89 | | | | -0.61 | | | | 0.59 | | | | 0.98 | | | | | | | | | | | | | | | | | | | | | | |
| **135-DR_0-90** | | | | | | | | | | | | | | | | | | | | | | | | | -0.88 | | | -0.75 | | | | -0.76 | | | | 0.75 | | | | 0.88 | | | | | | | | | | | | | | | | | | | | |
| **30-AR_0-90** | | | | | | | | | | | -0.29 | | | | | | | | | | | | | | | | | | | | | | | | | | | | | | | | | | | | | | | | | | | | | | | | | |
| **150-AL_0-90** | | | | | | | | | | | | -0.07 | | | | | | | | | | | | | | | | | | | | | | | | | | | | | | | | | | | | | | | | | | | | | | | | |
| **30-DL_0-90** | | | | | | | | | | | | | | | | | | | | | | | | | | | | | | | | | | | | | | | | | | | | | | | | | | | | | | | | | | | | |
| **150-DR_0-90** | | | | | | | | | | | | | | | | | | | | | | | | | | | | | | | | | | | | | | | | | | | | | | | | | | | | | | | | | | | | |
| **90-AU_45-135** | | | | | | | | | | | | | | 0.10 | | | 0.24 | | | | 0.76 | | | | | | -0.85 | | -0.37 | | -0.63 | | | | | | | | | | | | | | | | | | | | | | | | | | | | | |
| **90-DU_45-135** | | | | | | | | | | | | | | | | | | 0.33 | | | | 0.87 | | | -0.83 | | | | | -0.47 | | -0.51 | | | | | | | | | | | | | | | | | | | | | | | | | | | | |
| **0-UR_45-135** | | | | | | | | | | | | | | | | -0.19 | | | 0.53 | | | | 0.80 | | | | | -0.68 | -0.42 | | | -0.47 | | | | | | | | | | | | | | | | | | | | | | | | | | | | |
| **0-UL_45-135** | | | | | | | | | | | | | | | | | | | | 0.50 | | | | 0.77 | | -0.68 | | | | -0.62 | -0.52 | | | | | | | | | | | | | | | | | | | | | | | | | | | | | |
| **90-AU_30-150** | | | | | | | | | | | | | | | | | | 0.09 | | | 0.60 | | | | | | -1.04 | | | | | | 0.89 | | 0.64 | | | | | | | | | | | | | | | | | | | | | | | | | |
| **90-DU_30-150** | | | | | | | | | | | | | | | | | | | | | | 0.57 | | | -0.93 | | | | | | | | | 0.75 | | 0.80 | | | | | | | | | | | | | | | | | | | | | | | | |
| **0-UR_30-150** | | | | | | | | | | | | | | | | | | | | -0.24 | | | 0.60 | | | | | -0.95 | | | | | 0.86 | | | 0.82 | | | | | | | | | | | | | | | | | | | | | | | | |
| **0-UL_30-150** | | | | | | | | | | | | | | | | | | | | | | | | 0.58 | | -0.93 | | | | | | | | 0.79 | 0.90 | | | | | | | | | | | | | | | | | | | | | | | | | |
| **90-AU_60-120** | | | | | | | | | | | | | | | | | | | | | | -0.03 | | | | | -1.02 | | | | | | | | | | 0.92 | | 0.67 | | | | | | | | | | | | | | | | | | | | | |
| **90-DU_60-120** | | | | | | | | | | | | | | | | | | | | | | | | | -1.04 | | | | | | | | | | | | | 0.85 | | 0.85 | | | | | | | | | | | | | | | | | | | | |
| **0-UR_60-120** | | | | | | | | | | | | | | | | | | | | | | | | -0.19 | | | | -0.92 | | | | | | | | | 0.93 | | | 0.75 | | | | | | | | | | | | | | | | | | | | |
| **0-UL_60-120** | | | | | | | | | | | | | | | | | | | | | | | | | | -0.89 | | | | | | | | | | | | 0.88 | 0.88 | | | | | | | | | | | | | | | | | | | | | |
| **90-DU_0-90** | | | | | | | | | | | | | | | | | | | | | | | | | | | 0.07 | | | | | | | | | | | | | | | | | | | | | | | | | | | | | | | | | |
| **0-UL_0-90** | | | | | | | | | | | | | | | | | | | | | | | | | | | | 0.19 | | | | | | | | | | | | | | | | | | | | | | | | | | | | | | | | |
| **90-AU_0-90** | | | | | | | | | | | | | | | | | | | | | | | | | | | | | | | | | | | | | | | | | | | | | | | | | | | | | | | | | | | | |
| **0-UR_0-90** | | | | | | | | | | | | | | | | | | | | | | | | | | | | | | | | | | | | | | | | | | | | | | | | | | | | | | | | | | | | |

| | | | | | | | | | | | | | | | | | | | | | | | | | | | | | | | | | | | | | | | | | | | | | | | | | | | | | | | | | | | | |
|---|---|---|---|---|---|---|---|---|---|---|---|---|---|---|---|---|---|---|---|---|---|---|---|---|---|---|---|---|---|---|---|---|---|---|---|---|---|---|---|---|---|---|---|---|---|---|---|---|---|---|---|---|---|---|---|---|---|---|---|---|
|45-AR_45-135||||||||||||||||||||||||||||||-0.07|-0.18||1.09||||1.01||||||||||||||||||||||||
|45-DL_45-135||||||||||||||||||||||||||||||||0.03||1.03||||1.01|||||||||||||||||||||||
|135-AL_45-135||||||||||||||||||||||||||||||||0.03|||1.09||||1.04||||||||||||||||||||||
|135-DR_45-135||||||||||||||||||||||||||||||||||||1.04||||1.04|||||||||||||||||||||
|45-AR_105-165||||||||||||||||||||||||||||||||||-0.10|-0.12||0.82||||||||||||||||||||||||
|45-DL_105-165||||||||||||||||||||||||||||||||||||0.03||0.66|||||||||||||||||||||||
|135-AL_105-165||||||||||||||||||||||||||||||||||||-0.02|||0.67||||||||||||||||||||||
|135-DR_105-165||||||||||||||||||||||||||||||||||||||||0.62|||||||||||||||||||||
|45-AR_15-75||||||||||||||||||||||||||||||||||||||-0.46|-0.23||||||||||||||||||||||
|45-DL_15-75||||||||||||||||||||||||||||||||||||||||0.00|||||||||||||||||||||
|135-AL_15-75||||||||||||||||||||||||||||||||||||||||0.08|||||||||||||||||||||
|135-DR_15-75|||||||||||||||||||||||||||||||||||||||||||||||||||||||||||||
|30-UU_0-90|||||||||||||||||||||||||||||||||||||||||||||||||||||||||||||
|45-UU_0-90||||||||||||||||||||||||||||||||||||||||||||-0.11|||-1.03|-0.94||0.71|||-0.88|-0.84|-0.57|-0.51|0.98||||
|120-UU_0-90|||||||||||||||||||||||||||||||||||||||||||||||||||||||||||||
|135-UU_0-90|||||||||||||||||||||||||||||||||||||||||||||||-0.95|-0.90|0.80||||-0.89|-0.85|-0.54|-0.56||1.04|||
|150-UU_0-90|||||||||||||||||||||||||||||||||||||||||||||||||||||||||||||
|60-UU_0-90|||||||||||||||||||||||||||||||||||||||||||||||||||||||||||||
|0-UU_0-90||||||||||||||||||||||||||||||||||||||||||||||||0.10|||0.87||0.63|0.62|0.78|0.75|||1.02||
|90-UU_0-90||||||||||||||||||||||||||||||||||||||||||||||||||||1.01|0.61|0.71|0.69|0.81||||1.04|
|135-UU_105-165||||||||||||||||||||||||||||||||||||||||||||||||||-0.17|-0.98|-0.73||-1.10||||0.62|||
|45-UU_105-165|||||||||||||||||||||||||||||||||||||||||||||||||||-0.90|-0.78|-1.13||||0.56||||
|0-UU_30-150||||||||||||||||||||||||||||||||||||||||||||||||||||0.18|||-0.54||||0.52||
|90-UU_30-150||||||||||||||||||||||||||||||||||||||||||||||||||||||||-0.36||||0.53|
|45-UU_45-135||||||||||||||||||||||||||||||||||||||||||||||||||||||-0.03|0.26|0.64|1.18||||
|135-UU_45-135|||||||||||||||||||||||||||||||||||||||||||||||||||||||0.44|0.63||1.11|||
|0-UU_45-135||||||||||||||||||||||||||||||||||||||||||||||||||||||||0.17|||0.76||
|90-UU_45-135||||||||||||||||||||||||||||||||||||||||||||||||||||||||||||0.83|
|45-UU_15-75||||||||||||||||||||||||||||||||||||||||||||||||||||||||||-0.08|-0.94|-0.81|
|135-UU_15-75|||||||||||||||||||||||||||||||||||||||||||||||||||||||||||-0.95|-0.91|
|0-UU_60-120||||||||||||||||||||||||||||||||||||||||||||||||||||||||||||0.12|
|90-UU_60-120|||||||||||||||||||||||||||||||||||||||||||||||||||||||||||||

### Supporting text for Methods and Materials

All experiments were approved by the local ethics committee ("Commissione per l'Etica della Ricerca", University of Florence, January 17, 2024, No. 292) and conducted in accordance with the Declaration of Helsinki. Participants in all three experiments were naïve to the purpose of the study, had normal or corrected-to-normal vision, had no history of neurological conditions or brain injury, and were free of cognitive disorders. All participants provided informed consent prior to participation, and no monetary compensation was provided.

Main statistical analyses were conducted in R (version 4.4.2) and MATLAB (R2023b). Some additional descriptive analyses and assumption checks were performed using JASP (version 0.16.4).

#### Experiment 1: Rating dynamism of diagonals in still images

##### Participants

Two hundred and fifty adults took part in the study (F = 166, M = 82, Prefer not to disclose = 2; mean age = 25.4, SD = 9.6). 8% of the sample had a middle-school diploma, 68% had a high-school diploma, 17% held a bachelor's degree, 6% had a master's degree, and 1% held a PhD.

##### Set up

The study was conducted online using Google Forms. Participants were instructed to complete the experiment on a device with a sufficiently large display; however, no restrictions were imposed, and participants were asked to report the type of device they used. In total, 125 participants completed the task on a computer, 101 on a smartphone, and 24 on a tablet. Stimuli were generated using MATLAB R2018b and presented embedded in the online form. Responses were recorded automatically by Google Forms and later exported for analysis.

##### Stimuli

We created a set of 60 artificial stimuli, each composed of four equilateral quadrilaterals that were either equiangular squares or non-equiangular rhombuses, presented with the edges in cardinal or oblique orientations. These elements were arranged in linear configurations with vertical, horizontal, or oblique orientations. We refer to the angle with respect to the horizontal of the lines forming each element as *local* orientation, and to the layout axis of the four-element group as *global* orientation. In each stimulus, local and global orientations could be either congruent or incongruent, e.g., rhombuses with oblique 30°–150° edges could be arranged in a horizontal global orientation. Twenty stimuli contained elements of equal size (**Fig. 2A**), whereas the remaining 40 contained systematically varying element sizes along the configuration axis, forming wedge-shaped configurations (**Fig. 2B**). Element size manipulation allowed us to assess two potentially independent sources of perceived dynamism: configuration-based dynamism, arising from the spatial orientation of the composition, and element-based dynamism, produced by size gradients that may suggest depth or directional motion along the configuration axis.

Stimuli were designated using the convention "Global orientation–Size pattern_Local edge orientation". The first term specifies one of eight global configuration orientations (0°, 30°, 45°, 60°, 90°, 120°, 135°, and 150°). The two-letter size code describes the direction of element-size variation by combining three vertical components (Ascending [A], Descending [D], Uniform [U]) with three horizontal components (Right [R], Left [L], Uniform [U]), yielding the combinations AR, AL, DR, DL, AU, DU, UR, UL, and UU. "Ascending" and "Descending" indicate whether element size increased upward or downward along the vertical axis, whereas "Right" and "Left" indicate whether size increased toward the right or toward the left along the horizontal axis. All equal-size stimuli were designated "UU", as projections along both vertical and horizontal directions were labeled as Uniform. The final term specifies one of six local edge-orientation pairs (0°–90°, 45°–135°, 30°–150°, 60°–120°, 15°–75°, and 105°–165°). The first two local orientation pairs correspond to square elements, which have 90° internal angles and fourfold rotational symmetry. The remaining four correspond to rhombus-shaped elements with 30°–150° or 60°–120° internal angles and twofold symmetry, rotated so that either the short or the long diagonal aligns with the global configuration direction. As an example, the designation 0-UL_45-135 identifies a horizontal configuration (0°) in which element size is uniform along the vertical axis (U) and progressively increases from right to left along the horizontal axis (L), with 45°–135° local edges.

All stimuli were 900 × 900-pixel images. The equal-size square elements each measured 103 × 103 pixels, corresponding to an area of 10,609 pixels. In the wedge-shape conditions, each element was scaled to be approximately 30% smaller than the preceding one (e.g., in the square version, edge lengths were 152, 106, 74, and 52 pixels, yielding areas of 23,104, 11,236, 5,476, and 2,704 pixels, respectively). For rhombus elements, edge lengths were computed so that the resulting shapes matched the areas of the corresponding squares, with a maximum deviation of ~0.01% in the number of black pixels. In other words, the overall number of black pixels was tightly matched across all configurations, yielding comparable luminance levels.

The spacing between adjacent elements was fixed at 10 pixels in all conditions, both for the gap between facing edges and the distance between nearest vertices. This resulted in configurations with different overall spatial dispersion – that is, some arrangements appeared more compact, whereas others were more widely spaced (see **Fig. 2**). Elements were rendered in black on a uniform mid-gray background.

**Procedure**

After reading the instructions, participants proceeded through the questionnaire one stimulus at a time. On each page, a single stimulus was displayed together with three 5-point rating scales (−2 to +2) assessing perceived dynamism, tridimensionality, and direction. The labels associated with each numerical value were shown explicitly and remained visible throughout the task. For dynamism, –2 corresponded to completely static, –1 to fairly static, 0 to I don't know, +1 to fairly dynamic, and +2 to completely dynamic. For tridimensionality, –2 indicated completely flat, –1 fairly flat, 0 I don't know, +1 fairly tridimensional, and +2 completely tridimensional. For direction, –2 indicated completely raising, –1 fairly raising, 0 neither raising nor falling, +1 fairly falling, and +2 completely falling. Participants were required to respond to all three scales before advancing to the next page, where the following stimulus was presented.

To reduce response bias, the orientation of the numerical scale was randomized across participants: for 111 participants, –2 corresponded to "completely static", whereas for the remaining 139 participants –2 corresponded to "completely dynamic" (and similarly for the other two scales).

The experiment consisted of two blocks. The first block presented the 20 equal-size stimuli, followed by a second block with the 40 wedge-shaped stimuli. This block order was chosen to minimize potential carryover effects: because variations in element size may produce stronger impressions of motion, presenting size-varying stimuli first could plausibly have biased participants to judge the equal-size configurations as less dynamic. Within each block, the order of stimulus presentation was randomized.

#### Data processing and statistical analysis

For participants who received the reversed scale (i.e., –2 corresponding to "completely dynamic" and +2 to "completely static"), responses were inverted so that, for all participants, higher values consistently reflected higher perceived dynamism (and analogously for the other two scales). After recoding, responses were pooled across participants to obtain the distribution of ratings for each stimulus (**Fig. 3A**).

For each stimulus, we computed the mean rating for Dynamism, Tridimensionality, and Direction. We also computed the absolute value of the Direction rating as an index of perceived stability/instability, where 0 indicates perceived stability and non-zero values indicate perceived instability. The relationships among these perceptual measures were assessed by Pearson correlations computed on the stimulus-wise mean scores (**Fig. 3B**).

To summarize the effects of global and local orientation on perceived dynamism, stimuli were grouped into four categories obtained by crossing local edge orientation (cardinal vs. oblique) and global configuration orientation (cardinal vs. oblique). For each participant, dynamism ratings were averaged across all stimuli belonging to each category, and these participant averages were subsequently averaged across observers to obtain the group means for the four categories (**Fig. 3C**).

To assess the specific stimulus manipulations contributing to the observed effects, we conducted a series of contrasts between the dynamism ratings of targeted stimuli (**Table S1**). For each contrast, ratings were averaged within participants and then compared using both parametric (paired-samples t-tests) and non-parametric (Wilcoxon signed-rank tests) statistics. Because the rating data were ordinal and deviated from normality, statistical significance was evaluated on the basis of the Wilcoxon signed-rank tests, whereas t-statistics are reported as descriptive measures of effect direction and relative magnitude.

To complement mean-based analyses, we also performed chi-square tests comparing the full distributions of categorical responses across stimulus groups. This distribution-level assessment is well-suited to ordinal data and allows the detection of differences that may not be reflected solely in mean ratings.

### Experiment 2: Paired comparison rankings and a perceptual scale for perceived dynamism

#### Participants

Fifty adults took part in the study (F = 27, M = 23; mean age = 25.2, SD = 6.9). 70% of the sample had a high-school diploma, 20% held a bachelor's degree, 10% had a master's degree.

#### Set up

The experiment was conducted on a 24-inch Apple iMac (2021; macOS Sonoma 14.1) using custom Python scripts implemented in the open-source software OpenSesame (v4.0.24). Stimuli were displayed on the built-in Retina display (53.5 × 30.1 cm) at a resolution of 2240 × 1260 pixels. Participants viewed the display from a distance of 57 cm and responded using a computer mouse.

#### Stimuli

From the stimulus set described in Experiment 1, we constructed 168 distinct stimulus pairs designed to isolate specific visual factors hypothesized to influence the perception of dynamism. The pairs were grouped into four subsets, each aimed at testing a different visual factor by varying one parameter while holding the others constant.

The first subset tested the effect of global configuration orientation (64 pairs; **Table S2**). In these comparisons, stimuli differed in the orientation of the global configuration (0°, 45°, 90°, or 135°) while sharing the same local edge orientation (either cardinal or oblique) and the same pattern of element size variation. This allowed us to compare perceived dynamism across different global orientations while keeping both the local structure and the size gradient constant.

The second subset tested the effect of local edge orientation (72 pairs; **Table S3**). In these comparisons, stimuli differed in the orientation of the local edges (0°–90°, 45°–135°, 30°–150°, 60°–120°, 15°–75°, or 105°–165°) while sharing the same global configuration orientation and the same pattern of element size variation. This allowed us to compare perceived dynamism across different local edge structures while keeping both the global configuration and the size gradient constant.

The third subset tested the effect of element size variation (20 pairs; **Table S4**). In these comparisons, stimuli shared the same global configuration orientation and the same local edge orientation but differed in the direction of the element size gradient (A-R vs D-L, A-L vs D-R, A-U vs D-U, or U-R vs U-L). This allowed us to compare perceived dynamism across different size-variation patterns while keeping both the global configuration and the local edge structure constant.

Finally, a fourth subset examined the interaction between global configuration orientation and local edge orientation (12 pairs; **Table S5**). In these comparisons, stimuli combined cardinal or oblique global configurations (0°, 90° vs 45°, 135°) with cardinal or oblique local edge orientations (0°–90° vs 45°–135°), while sharing the same pattern of element size variation. This allowed us to compare configurations in which global and local orientations were either aligned (both cardinal or both oblique) or misaligned (cardinal–oblique combinations), thereby assessing how the relationship between global and local orientation structures modulates perceived dynamism.

Pairs of configurations were presented against a uniform gray background and were scaled with OpenSesame to 40% of their original size, subtending approximately 8.6° × 8.6° of visual angle at a viewing distance of 57 cm. All tested stimulus pairs are illustrated in Supplementary Folder 1.

**Procedure**

Participants performed a pairwise perceptual judgment task in which they compared two configurations presented simultaneously and indicated which image appeared more dynamic. Responses were made using the mouse by clicking on a five-point response scale ranging from −2 to +2, where −2 indicated that the left image was definitely more dynamic, −1 that the left image was probably more dynamic, 0 that the two images were perceived as equally dynamic, +1 that the right image was probably more dynamic, and +2 that the right image was definitely more dynamic.

On each trial, two configurations were displayed symmetrically to the left and right of screen center, with their centers positioned at an eccentricity of 7.2° of visual angle. The response scale was presented below the stimuli, with the central value (0) positioned midway between the two images, while the extreme values (−2 and +2) were placed below the left and right stimuli, respectively. The two images remained on the screen until a response was made.

The experiment consisted of 1008 trials, corresponding to 168 stimulus pairs presented six times each. Within each pair, the left–right position of the two stimuli was counterbalanced across repetitions (three trials per spatial arrangement). Trials were presented in random order. To reduce fatigue, trials were divided into six blocks separated by short self-paced breaks.

#### Data processing and statistical analysis

For each stimulus pair, responses were averaged across participants to obtain a mean relative dynamism score (**Table S6**). Consistency between Experiments 1 and 2 was assessed by correlating these scores across pairs with the corresponding differences in mean dynamism ratings obtained in Experiment 1 (**Table S7**). Stimulus pairs were grouped according to the specific visual factor they were designed to probe — global configuration orientation, local edge orientation, the interaction between global and local orientations, and the direction of element size variation — and correlations were computed separately for each subset (**Fig. 4A**) using Spearman's rank correlation coefficient ($\varrho$). The coefficient of determination ($R^2$), derived from linear fits, is reported to facilitate interpretation of the strength of the relationship.

Subsequently, we focused on a subset of eight stimuli for which all possible pairwise combinations were tested (28 pairs in total; see **Fig. 4B**). This complete set of comparisons allowed us to derive a perceptual scale of perceived dynamism. To this end, we first computed response matrices separately for neutral responses (0), intermediate responses (–1 and +1), and extreme responses (–2 and +2). Because extreme responses were relatively infrequent, intermediate and extreme responses were combined to obtain a binarized preference matrix, in which we grouped responses favoring the left stimulus (–2 and –1) and those favoring the right stimulus (+1 and +2). The resulting matrix contained, for each stimulus pair, the proportion of trials in which one stimulus was judged as more dynamic than the other (**Fig. 4B**). This binarized comparison matrix was then used as input to an extended maximum-likelihood implementation of Thurstone's scaling method (11), yielding a quantitative perceptual scale of perceived dynamism (**Fig. 4C**). In this scale, each stimulus is assigned a position along a latent psychological dimension reflecting its perceived dynamism, with higher values indicating stimuli more frequently judged as more dynamic. Differences between scale values reflect the relative perceptual distances between stimuli.

Finally, to assess the correspondence between the perceptual scale derived from pairwise comparisons and the ratings obtained in Experiment 1, we computed a Spearman correlation between the scale values and the mean dynamism ratings from the questionnaire (**Fig. 4D**). This analysis provided a cross-validation of the two measurement approaches. The coefficient of determination ($R^2$), derived from linear fits, is reported to facilitate interpretation of the strength of the relationship.

### Experiment 3: Pupil responses to perceived dynamism

#### Participants

Thirty-five adults took part in the study (F = 23, M = 12; mean age = 23.08, SD = 2.44). Twenty-nine participants had a high-school diploma, four held a bachelor's degree, and one had a middle-school diploma. None of the participants reported taking medication at the time of the experiment.

#### Set up

The experiment took place in a dimly lit, quiet room. Participants were seated comfortably with their head stabilized using a chinrest positioned 57 cm from the display. Eye movements and pupil size were recorded using a remote infrared eye tracker (EyeLink 1000, SR Research Ltd.) positioned below the monitor. Stimuli were presented on a gamma-calibrated AOC 27G2U/BK monitor (resolution: 1920 × 1080 pixels; refresh rate: 120 Hz), subtending approximately 61° × 34° of visual angle at the viewing distance. The session began with a standard 9-point calibration procedure for the eye-tracker, after which eye-tracking data were continuously recorded at a sampling rate of 500 Hz. Stimulus presentation and response collection were controlled using the open-source software OpenSesame (4.0.24).

#### Stimuli

To select the stimulus sample, the 60 images used in Experiment 1 were ordered according to their mean dynamism ratings (**Fig. 3A**). Based on these ratings, we identified the 12 most static and the 12 most dynamic configurations. From this pool, we selected a final subset of 13 stimuli (see **Fig. 5B**). This subset included (i) four static stimuli in which local and global orientations were congruent: two stimuli combined cardinal local edges (0°–90°) with cardinal global configurations (0° and 90°), and two combined oblique local edges (45°–135°) with oblique global configurations (45° and 135°); (ii) four static stimuli in which local and global orientations were incongruent: two stimuli combined oblique local edges (45°–135°) with cardinal global configurations (0° and 90°), and two combined cardinal local edges (0°–90°) with oblique global configurations (45° and 135°); (iii) four dynamic stimuli with wedge-shaped elements, covering the same global orientations as the static set (0°, 90°, 45°, and 135°). All dynamic configurations contained oblique local edge orientations (30°–150° for cardinal global orientations, and 105°–165° for oblique global orientations). It was not possible to construct an alternative dynamic set with cardinal local orientations, as all highly dynamic stimuli included oblique local features. These dynamic stimuli also conveyed clear directional properties along different axes (vertical, horizontal, leftward, or rightward), depending on their internal structure; and (iv) one additional dynamic stimulus with equal-size elements, that matched one of the dynamic wedge-shaped stimuli in both global orientation (135°) and local edge orientation (105°–165°), differing only in the absence of size variation across elements.

Stimuli were presented against a uniform gray background. A scrambled control stimulus was designed to have matched luminance and size, but no recognizable visual motif, which served as the baseline for measuring phasic changes in pupil size.

#### Procedure

The experiment consisted of two sequential phases. The first phase was designed to measure pupillary responses under controlled fixation conditions, whereas the second phase was designed to collect perceptual judgments for each stimulus.

The pupillometry phase consisted of 104 trials, corresponding to 13 stimuli presented eight times each in random order. Each trial began with the presentation of a scrambled baseline stimulus for 3–4 s (uniform jitter), followed by the target image displayed centrally for 6 s. Images subtended 7° of visual angle. This relatively small size was chosen to allow observers to perceive the entire configuration without the need for eye movements, which may bias the measurement of pupil size. Participants were instructed to maintain fixation throughout the trial while pupil size was continuously recorded. Trials were divided into three blocks separated by short self-paced breaks to reduce eye fatigue. Before resuming each block, eye-tracker calibration accuracy was verified.

The second phase consisted of a single block of 13 trials, in which each stimulus was presented once for 8 s at the center of the screen, subtending 21° of visual angle. Following each image, participants rated the configuration on the three perceptual dimensions used in Experiment 1 – dynamism, tridimensionality, and direction – using the same 5-point scale ranging from −2 to +2. Observers responded by clicking the desired option on the screen with the computer mouse.

**Data processing and statistical analysis**

Pupil data

Pupil size was recorded from the left eye, in arbitrary units of area. We retained only the values measured during fixation events, as identified by the eye tracker (i.e., measures during blinks and saccades were discarded). We then further removed values below 2.5 standard deviations from each participant's average pupil size and used a velocity-based criterion to remove implausibly fast changes in pupil size, as well as 120 ms of recordings around these values. These procedures were adopted to minimize the impact of artifacts such as partial occlusion of the pupil by the eyelid. Trials in which more than 40% of the data were missing were discarded. The gaps in the remaining trials were linearly interpolated, then traces were smoothed with cubic splines. Finally, traces were down-sampled to 20-ms epochs by taking the median pupil size within each time bin. To better cope with inter-individual differences, we z-transformed pupil size values separately for each participant. We then computed a trial-wise baseline level by taking the median pupil size between -250 and 250 ms, centered on stimulus presentation. Trials with extreme baseline values (exceeding +-2 standard deviations from the mean) were discarded; otherwise, pupil traces were realigned by subtraction to this baseline. The preprocessing pipeline largely follows previous studies and the default parameters of the R library Pupilla (1). Data cleaning procedures led to discarding 14% of trials per participant on average.

We submitted the remaining data to Principal Components Analysis (PCA), which recovered one dominant dilation trend across all trials (>80% explained variance). We thus used PC1 scores as a summary measure of pupillary response magnitude.

For each participant, PC1 scores were averaged across repetitions for each stimulus and submitted to a repeated-measures ANOVA with Configuration as the within-subject factor to assess whether pupillary responses differed across the 13 configurations. We then tested whether this variation reflected perceived dynamism rather than image identity per se by computing, for each participant, the Spearman correlation between the dynamism ratings assigned to each stimulus and the corresponding PC1 scores (**Fig. 5B**). Correlation coefficients were finally tested against zero using one-sample t-tests.
